# TGF-β Coordinates Alanine Synthesis and Import for Myofibroblast Differentiation in Pulmonary Fibrosis

**DOI:** 10.1101/2025.07.23.666333

**Authors:** Fei Li, Niv Vigder, David R Ziehr, Mari Kamiya, Hung N Nguyen, Matthew L Steinhauser, Edy Y Kim, William M Oldham

## Abstract

Idiopathic pulmonary fibrosis (IPF) is a progressive interstitial lung disease marked by aberrant fibroblast-to-myofibroblast differentiation, a process that requires metabolic reprogramming. We identify alanine as a critical metabolite that confers metabolic flexibility to support differentiation. TGF-β increases alanine by activating both its synthesis and import in normal and IPF lung fibroblasts. Alanine is synthesized primarily by GPT2, which is regulated by a glutamine–glutamate–α-ketoglutarate axis. Inhibiting GPT2 depletes alanine and suppresses TGF-β-induced expression of α-SMA and COL1A1, an effect reversed by alanine supplementation. We also identify SLC38A2 as a key transporter of both alanine and glutamine that is upregulated by TGF-β and alanine deprivation. Together, SLC38A2 and GPT2 activities converge to maintain intracellular alanine levels to support myofibroblast differentiation. Mechanistically, alanine deficiency suppresses glycolysis and depletes tricarboxylic acid cycle intermediates, while supplementation provides carbon and nitrogen for intracellular glutamate and proline biosynthesis, particularly in the absence of glutamine. Combined inhibition of GPT2 and SLC38A2 suppresses fibrogenic responses in fibroblasts and in human precision-cut lung slices, highlighting a potential therapeutic strategy for fibrotic lung disease.

## Introduction

Idiopathic pulmonary fibrosis (IPF) is a progressive and lethal lung disease with a median survival of 3-5 years following diagnosis 1. Although two antifibrotic drugs, pirfenidone and nintedanib, have been approved for IPF treatment, their efficacy is limited to slowing disease progression 2,3. Thus, a deeper understanding of underlying disease mechanisms is essential for the development of more effective antifibrotic therapies.

Fibroblast-to-myofibroblast differentiation is a key step in fibrosis progression 4. Under the influence of profibrotic factors such as TGF-β, pulmonary fibroblasts differentiate to myofibroblasts characterized by significant metabolic reprogramming, including increased nutrient uptake and activation of anabolic pathways that promote extracellular matrix deposition and contractile force generation 5. Recent studies have shown that myofibroblast differentiation requires changes in glucose and glutamine metabolic pathways 6. TGF-β promotes the uptake of extracellular glucose and its subsequent metabolism into glycine and lactate, both of which are building blocks for excessive collagen synthesis 7,8.

Similarly, glutamine metabolism contributes to the collagen deposition that drives pathological tissue remodeling in fibrosis 9–11. Glutamine is a vital nutrient for rapidly proliferating cells as it supports metabolic and biosynthetic reactions 12,13. Strategies targeting glutamine metabolism are emerging for the treatment of IPF, with glutaminase (GLS) identified as a potential therapeutic target for fibrosis 9,14. However, cancer-related studies have revealed that targeting glutamine metabolism showed limited efficacy in vivo due to metabolic adaptability that enables cells to bypass glutamine-dependent metabolic requirements for proliferation 15. This metabolic flexibility may involve compensatory pathways mediated by transaminases and alternative metabolites such as glutamate, asparagine, serine, α-ketoglutarate, and pyruvate 16–20. Additionally, while non-essential amino acid (NEAAs, e.g., glutamine, serine, and alanine) can be synthesized intracellularly, the tight regulation of their intracellular concentration depends largely on extracellular sources, particularly when synthesis is limited 21,22. Through import and export, the extracellular metabolite pool offers the buffering capacity that is required for cells to maintain a narrow, homeostatic range of metabolite concentrations intracellularly. The reliance on extracellular metabolite pools also suggests that intracellular synthesis alone is insufficient to meet the metabolic demands of activated fibroblasts, exemplified by the fact that targeted inhibition of the activity of key transaminases does not completely inhibit myofibroblast differentiation 6. Metabolic adaptability, characterized by flexible utilization of both intracellular and extracellular metabolite sources, may enable myofibroblasts to tolerate fluctuations in nutrient availability and likely limits the efficacy of metabolism-targeted antifibrotic therapies.

Building upon this hypothesis, we investigated whether and how specific non-essential amino acids (NEAAs) act as compensatory substrates during myofibroblast differentiation. We demonstrate that alanine provision, through de novo synthesis by glutamate-pyruvate transaminase 2 (GPT2) and import by solute carrier family 38 member 2 (SLC38A2), is essential for myofibroblast differentiation. Combined pharmacological inhibition of both alanine synthesis and uptake significantly suppresses this process, revealing a previously unrecognized and potentially effective therapeutic strategy for IPF.

## Results

### TGF-β-induced myofibroblast differentiation increases intracellular alanine

To examine how medium composition affects myofibroblast differentiation, we treated primary normal human lung fibroblasts (NHLF) with TGF-β (2 ng/ml) in one of two commonly-used fibroblast culture media – Dulbecco’s Modified Eagle Medium (DMEM) or Fibroblast Basal Medium (FBM). DMEM is a general-purpose medium containing glutamine and branched-chain amino acids (BCAAs; valine, leucine, and isoleucine), but lacking NEAAs (alanine, proline, glutamate, asparagine, and aspartate) (**Table 1**). FBM, a fibroblast-specific medium, has a more complex proprietary amino acid composition that contains NEAAs (**Fig. EV1A-E**). To assess the metabolic changes following TGF-β, we used liquid chromatography – high-resolution accurate mass spectrometry (LC-MS) to profile the intracellular metabolome of NHLF cells after 48 h of TGF-β treatment (**Fig. 1A**). Principal components analysis (PCA) of ∼100 measured metabolites revealed a clear separation among experimental groups, with the first principal component (50% of variance) driven by medium composition and the second principal component (27% of variance) reflecting the effects of TGF-β treatment (**Fig. 1B**).

**Table1.** Amino acid composition of DMEM.

| **Amino Acids (DMEM)** | **mM** |
| --- | --- |
| Alanine | Not Added |
| L-Arginine hydrochloride | 0.39810428 |
| Asparagine | Not Added |
| Aspartate | Not Added |
| L-Cystine 2HCl | 0.20127796 |
| Glutamate | Not Added |
| L-Glutamine | 4 |
| Glycine | 0.4 |
| L-Histidine hydrochloride-H<sub>2</sub>O | 0.2 |
| L-Isoleucine | 0.8015267 |
| L-Leucine | 0.8015267 |
| L-Lysine hydrochloride | 0.7978142 |
| L-Methionine | 0.20134228 |
| L-Phenylalanine | 0.4 |
| Proline | Not Added |
| L-Serine | 0.4 |
| L-Threonine | 0.79831934 |
| L-Tryptophan | 0.07843138 |
| L-Tyrosine disodium salt dihydrate | 0.39846742 |
| L-Valine | 0.8034188 |

**Figure 1.**
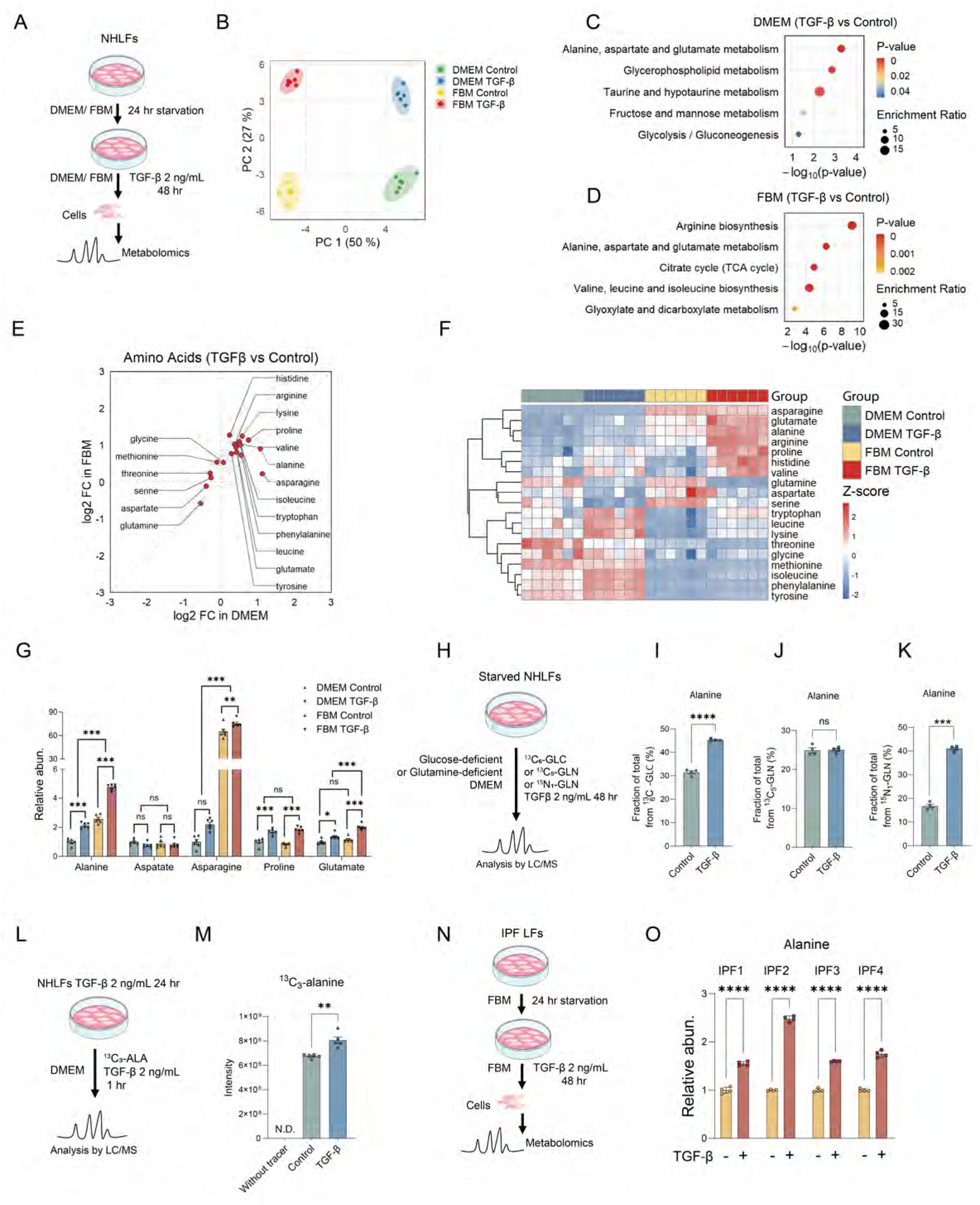
Increased intracellular alanine levels during myofibroblast differentiation. (A) Schematic overview of the experimental design for primary normal human lung fibroblasts (NHLFs) treated with TGF-β (2 ng/ml) in DMEM or FBM. Samples were collected and analyzed after 48 h using LC-MS (n = 6 for each condition). (B) PCA analysis of the experimental groups. (C, D) Enriched metabolite sets after 48 h TGF-β treatment in DMEM (C) and FBM (D), based on differential metabolites with FC ≥ 1.5 and P ≤ 0.05. Enrichment ratio = Hits / Expected. (E) Comparison of intracellular amino acid changes induced by TGF-β treatment in DMEM vs. FBM conditions. (F) Heatmap showing the intracellular levels of amino acids in fibroblasts cultured in DMEM or FBM with or without TGF-β stimulation for 48 h. Each column represents a sample, and each row corresponds to an amino acid. Colors indicate Z-score–normalized abundance, with red and blue representing higher and lower relative levels, respectively. (G) Relative quantification of the intracellular abundances of alanine, aspartate, asparagine, proline, and glutamate. (H) Schematic overview of LC-MS-based stable isotope tracing with [U-^13^C_6_]-glucose (^13^C_6_-GLC), [U-^13^C_5_]-glutamine (^13^C_5_-GLN), or [α-^15^N_1_]-glutamine (^15^N_1_-GLN) for 48 h. (I-K) Stable isotope label enrichment of alanine from ^13^C_6_-GLC (I), ^13^C_5_-GLN (J), or ^15^N_1_-GLN (K). (L) Schematic overview of short-term uptake assays using isotope tracing with [U-^13^C_3_]-alanine (^13^C_3_-ALA) for 1 h. (M) Signal intensity of intracellular ¹³C₃-alanine (undetectable without tracer). (N) Schematic overview of the experimental design for IPF human lung fibroblasts from four donors (IPF-LFs) treated with TGF-β (2 ng/ml) in FBM for 48 h. (O) Relative quantification of intracellular alanine. Each metabolite in G: one-way ANOVA; I, J, K, M, and O: unpaired t-test. Data are presented as mean ± SEM. ns, p > 0.05; *p < 0.05; **p < 0.01; ***p < 0.001; ****p < 0.0001.

Metabolite set enrichment analyses revealed an upregulation of amino acid metabolism following TGF-β treatment in DMEM and FBM (**Fig. 1C, D**). The concentrations of numerous amino acids were differentially regulated by TGF-β, regardless of culture medium. Notably, intracellular glutamine was decreased, whereas levels of several non-essential amino acids (NEAAs), including proline, alanine, and glutamate, were increased (**Fig. 1E**). Comparison of DMEM- and FBM-cultured cells revealed baseline and TGF-β–induced differences in metabolite profiles, highlighting that medium composition can shape cellular metabolism and influence responses to pro-fibrotic stimuli. (**Fig. 1F**). When cells are deprived of an exogenous supply of the five NEAAs (i.e., DMEM), TGF-β increased the intracellular concentrations of alanine, proline, and glutamate (**Fig. 1G**), suggesting enhanced intracellular synthesis. Among these, the intracellular concentration of alanine, irrespective of TGF-β treatment, was significantly higher in FBM-when compared with DMEM-cultured cells, suggesting that the intracellular level of alanine is regulated, at least in part, by the extracellular alanine pool (**Fig. 1G**).

Given the observed increases in alanine following TGF-β treatment, we next sought to determine the metabolic origins of alanine, as determined by LC-MS-based stable isotope tracing experiments using [U-^13^C_6_]-glucose, [U-^13^C_5_]-glutamine, or [α-^15^N_1_]-glutamine (**Fig. 1H**). [U-^13^C_6_]-glucose tracing confirmed that glucose serves as a key carbon contributor to alanine synthesis (**Fig. 1I**), consistent with previous studies 23. TGF-β treatment further increased the fraction of glucose-labeled alanine from ∼30% to ∼45% (**Fig. 1I**). We found that ∼25% and ∼20% of the total alanine pool was labeled from [U-^13^C_5_]-glutamine and [α-^15^N_1_]-glutamine, respectively, and that the TGF-β further increased the fraction of ^15^N-alanine labeling from ^15^N_1_-glutamine to ∼40% (**Fig. 1H, J, K**). Taken together, these data suggest that, apart from glucose, glutamine is a significant carbon and nitrogen source for alanine synthesis under both baseline conditions and TGF-β treatment.

To quantify alanine uptake, we conducted short-term (1 h) labeling experiments using [U-^13^C_3_]-alanine in control and TGF-β-pre-treated cells in DMEM (**Fig. 1L**). Labeled intracellular alanine were detected, with TGF-β further enhancing its accumulation, indicating increased alanine uptake under both baseline and TGF-β treatment conditions (**Fig. 1M**).

To determine whether these perturbations in alanine metabolism occur in a disease context, we cultured primary fibroblasts isolated from four IPF patients (IPF1-4) in FBM, and confirmed that TGF-β significantly increased alanine levels intracellularly by ∼1.5-2.5 fold (**Fig. 1N, O**), consistent with previous studies reporting higher alanine levels in lungs of IPF patients 24,25. These results suggest that TGF-β increases intracellular alanine during myofibroblast differentiation and pulmonary fibrosis.

### Glutamine promotes TGF-β-induced GPT2 expression to support myofibroblast differentiation

Alanine synthesis is mediated by glutamate-pyruvate transaminases (GPTs). In this process, glutamine is converted to glutamate by glutaminase (GLS), and glutamate subsequently undergoes transamination with pyruvate by GPT1/2 to generate alanine and α-KG (**Fig. 2A**). Our [α-^15^N_1_]-glutamine tracing data demonstrated a 2.5-fold increase in transaminase activity following TGF-β (**Fig. 1K**). We assessed the protein expression of cytosolic GPT1 and mitochondrial GPT2 during TGF-β-induced myofibroblast differentiation in DMEM and found that TGF-β increased GPT2 expression but had no effect on GPT1 (**Fig. 2B-E**).

**Figure 2.**
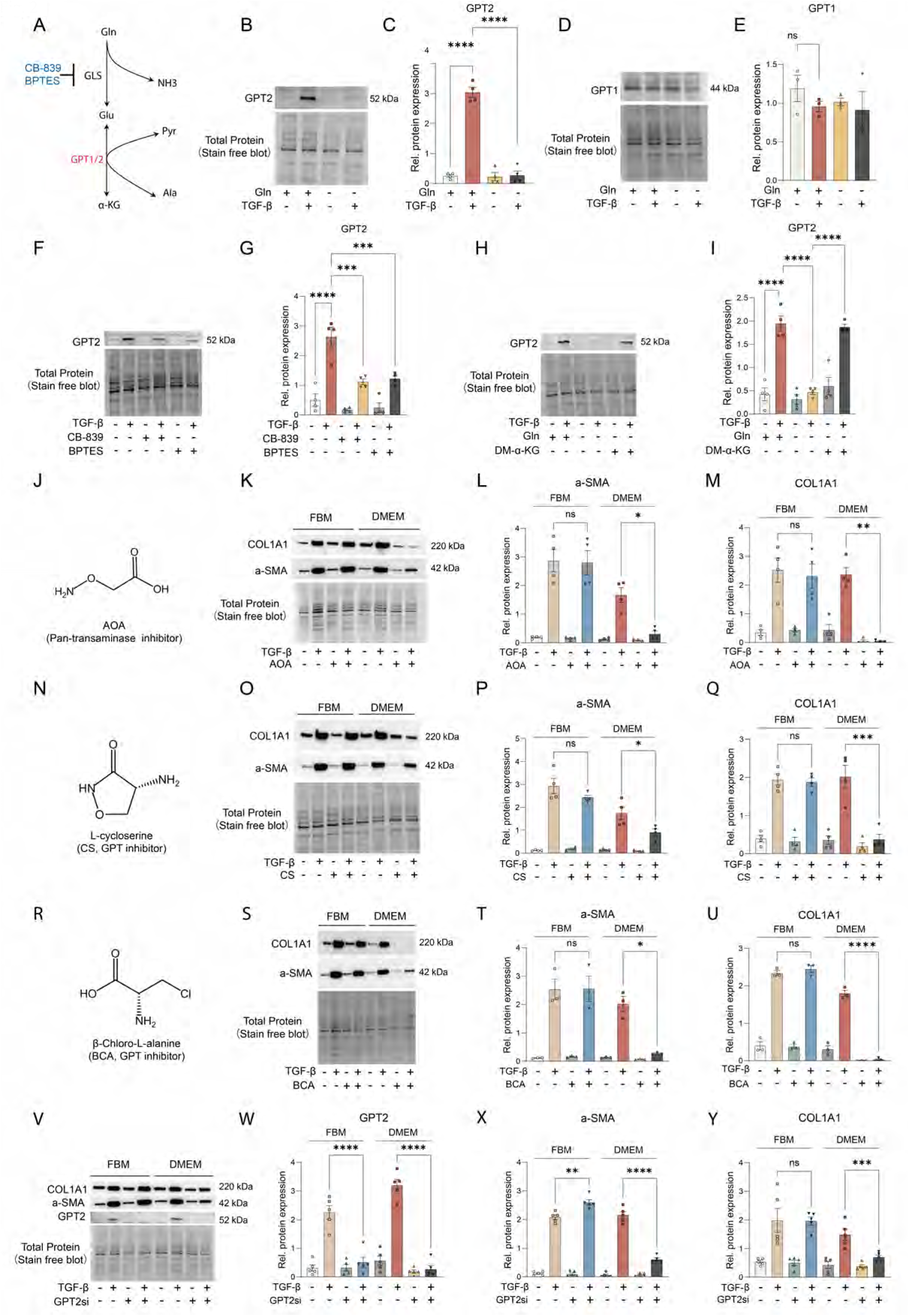
Glutamine promotes TGF-β-induced GPT2 protein expression to support myofibroblast differentiation. (A) Schematic overview of glutamine catabolism and alanine synthesis involving GPT1 and GPT2. (B, C) Western blot analysis and quantification of GPT2 protein expression in NHLFs treated with TGF-β in glutamine-deficient or glutamine-sufficient (2 mM) DMEM for 48 h. (D, E) Western blot analysis and quantification of GPT1 protein expression in NHLFs treated with TGF-β in glutamine-deficient or glutamine-sufficient (2 mM) DMEM for 48 h. (F, G) Western blot analysis and quantification of GPT2 in NHLFs following stimulation with TGF-β and treatment with GLS-1 inhibitors CB-839 (10 µM) or BPTES (5 µM) in DMEM for 48 h. (H, I) Western blot analysis and quantification of GPT2 in NHLFs following stimulation with TGF-β in glutamine-deficient or glutamine-sufficient DMEM for 48 h, with or without supplementation with dimethyl-α-ketoglutarate (DM-αKG, 5 mM). (J) Chemical structure of the pan-transaminase inhibitor AOA. (K-M) NHLF cells were stimulated with TGF-β in FBM or DMEM, with or without AOA (1 mM) for 48 h. α-SMA (L) and COL1A1 (M) expression were examined by Western blot. (N) Chemical structure of the GPT1/2 inhibitor CS. (O-Q) NHLF cells were stimulated with TGF-β in FBM or DMEM, with or without CS (100 µM) treatment for 48 h. α-SMA (P) and COL1A1 (Q) expression were examined by Western blot. (R) Chemical structure of the GPT1/2 inhibitor BCA. (S-U) NHLF cells were stimulated with TGF-β in FBM or DMEM, with or without BCA (100 µM) treatment for 48 h. α-SMA (T) and COL1A1 (U) expression were examined by Western blot. (V-Y) NHLF cells were transfected with control siRNA or GPT2 siRNA for 24 h, followed by 24-h starvation. Cells were then stimulated with TGF-β in FBM or DMEM for 48 h. GPT2-knockdown efficiency (W), α-SMA (X) and COL1A1 (Y) expression were examined by Western blot. Individual data points represent biological replicates. C, E, G, and I: one-way ANOVA vs. TGF-β; L, M, P, Q, W–Y: one-way ANOVA vs. TGF-β within each medium. ns, p > 0.05; *p < 0.05; **p < 0.01; ****p < 0.001; ****p < 0.0001.

To determine the role of GPT2 in myofibroblast differentiation, we first examined the effects of TGF-β in medium lacking glutamine (Gln), a primary source of glutamate for GPT2-catalyzed transamination of pyruvate to alanine. Interestingly, TGF-β did not increase GPT2 expression in the absence of glutamine (**Fig. 2B, C**). Similarly, the GLS-1 inhibitors telaglenastat (CB-839) and BPTES suppressed the TGF-β-induced upregulation of GPT2 protein expression in glutamine-containing medium (**Fig. 2F, G**), despite discrepancies with previous studies 16,26. Glutamine deficiency may decrease protein expression by impairing their translation or stability, a process that can be rescued by α-KG 27. Indeed, we found that supplementation with a cell-permeant analogue of α-KG [dimethyl (DM)-α-KG] enabled the TGF-β-induced upregulation of GPT2 protein expression even in the absence of glutamine (**Fig. 2H, I**). Taken together, these data suggest that a glutamine–glutamate–α-ketoglutarate (αKG) metabolic axis promotes TGF-β–induced upregulation of GPT2.

We next examined direct inhibition of GPT2 by small molecules to investigate its role in TGF-β-induced myofibroblast differentiation. We first used the pan-transaminase inhibitor, aminooxyacetic acid (AOA) 28, which has been shown previously to block α-SMA and collagen production during myofibroblast differentiation 10. Unexpectedly, AOA did not inhibit TGF-β-mediated upregulation of α-SMA and COL1A1 in FBM, although it was effective in DMEM as previously demonstrated (**Fig. 2J-M**) 10. We then tested the effects of the GPT1/2 inhibitors, L-cycloserine (CS) and β-chloro-L-alanine (BCA) 29. Like AOA, both CS and BCA suppressed TGF-β–induced upregulation of α-SMA and COL1A1 protein expression at lower concentrations (100 µM) in DMEM, but had no effect in FBM (**Fig. 2N-U, Fig. EV2A–D**). Given that CS and BCA lack selectivity for the individual isoforms of GPT (*i.e.,* GPT1 *vs.* GPT2), we next sought to study the role of each isoform separately using RNA interference. GPT1 knockdown had no effect on myofibroblast differentiation in either FBM or DMEM, as assessed by the relative protein expression of COL1A1 and α-SMA (**Fig. EV2E-H**). By contrast, GPT2 knockdown significantly decreased the TGF-β-induced upregulation of α-SMA and COL1A1 protein expressions in DMEM (**Fig. 2V-Y**). GPT2 knockdown did not affect the activation of the Smad3 signaling pathway following shorter TGF-β treatment (**Fig. EV2I, J**). Together, these results suggest that GPT2 is required for myofibroblast differentiation only under NEAA-deficient conditions (*i.e.,* DMEM). In contrast, when exogenous NEAAs are available (*i.e.,* FBM), GPT2 is redundant with respect to TGF-β-induced myofibroblast differentiation.

### Alanine is essential for α-SMA expression and collagen production in myofibroblasts

Next, we investigated whether the role of GPT2 in supporting myofibroblast differentiation depends on alanine metabolism. To this end, we analyzed intracellular metabolite profiles following either genetic knockdown or pharmacological inhibition of GPT2 in DMEM-cultured fibroblasts. In the absence of TGF-β, AOA and BCA treatment significantly altered the intracellular concentration of 24 and 27 metabolites, respectively, while both CS and GPT2 knockdown separately led to differential regulation of 4 metabolites (**Fig. 3A**). Alanine was the only metabolite that was consistently downregulated in each of the four treatments (**Fig. 3A, B**). Notably, under TGF-β stimulation, alanine remained the most significantly altered amino acid across all treatment groups (**Fig. EV3A-D**).

**Figure 3.**
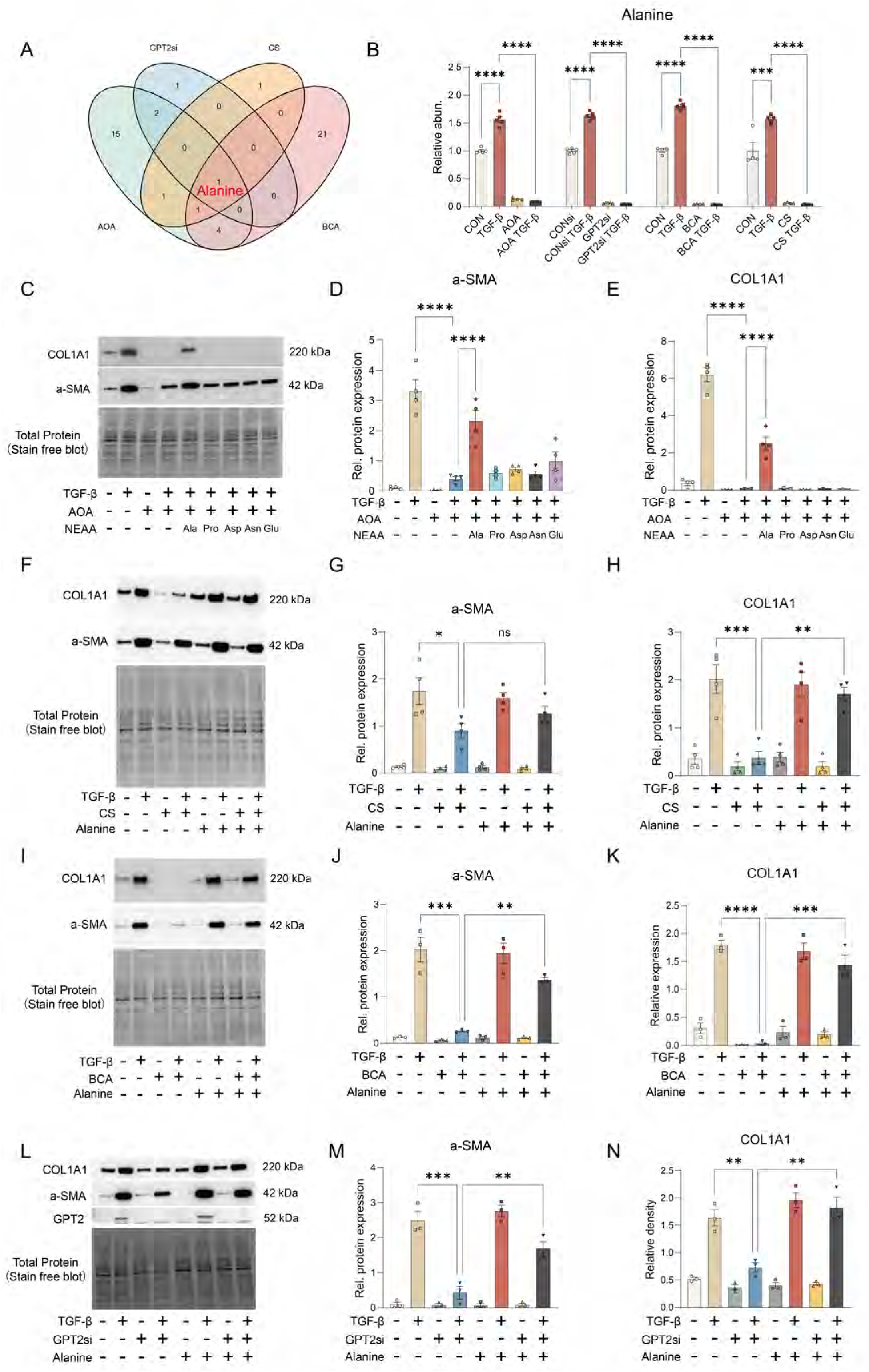
Alanine is essential for myofibroblast differentiation. (A) Venn diagram depicting the number of differentially regulated metabolites across four treatment conditions compared to their respective control groups, with alanine as the only shared metabolite. Differential metabolites: FC ≥ 1.5 and P ≤ 0.05. (B) Intracellular alanine levels in NHLF cells treated with TGF-β for 48 h, with or without GPT2 knockdown or inhibition in DMEM. (C-E) Western blot analysis (C) and quantification of α-SMA (D) and COL1A1 (E) expression in NHLF cells treated with TGF-β for 48 h in DMEM, with or without individual NEAA supplementation (2 mM each). (F-H) NHLF cells were stimulated with TGF-β in DMEM for 48 h, with or without CS (100 µM) and alanine supplementation (2 mM). α-SMA (G) and COL1A1 (H) expression were examined by Western blot. (I-K) NHLF cells were stimulated with TGF-β in DMEM for 48 h, with or without BCA (100 µM) and alanine supplementation (2 mM). α-SMA (J) and COL1A1 (K) expression were examined by Western blot. (L-N) NHLF cells were transfected with control siRNA or GPT2 siRNA for 24 h, followed by 24-h starvation. Cells were then stimulated with TGF-β in DMEM for 48 h, with or without alanine supplementation. α-SMA (M) and COL1A1 (N) expression were examined by Western blot. Individual data points represent biological replicates. B vs. TGF-β and D, E, G, H, J, K, M, and N vs. knock down/inhibitor plus TGF-β were analyzed using one-way ANOVA. Data are presented as mean ± SEM. ns, p > 0.05; *p < 0.05; **p < 0.01; ***p < 0.001; ****p < 0.0001.

To determine whether alanine deficiency impairs myofibroblast differentiation, we performed rescue experiments by supplementing 2 mM exogenous alanine, as per previously-used doses 23,30,31. Given that DMEM lacks five NEAAs, we compared the effects of alanine supplementation with those of the other four NEAAs. Supplementation with alanine – but not proline, aspartate, asparagine, or glutamate – significantly reversed the inhibitory effect of AOA on TGF-β-induced α-SMA and COL1A1 protein upregulation in DMEM (**Fig. 3C-E**). Similarly, alanine supplementation restored the TGF-β–induced upregulation of α-SMA and COL1A1 that was suppressed by CS (**Fig. 3F-H**), BCA (**Fig. 3I-K**), or GPT2 knockdown (**Fig. 3L-N**). These data suggest that alanine is required for myofibroblast differentiation and can bypass the anti-fibrotic effects of GPT2 inhibition.

### SLC38A2 facilitates alanine uptake to support myofibroblast differentiation

Given that inhibition of intracellular alanine synthesis did not impair myofibroblast differentiation in alanine-replete FBM, we hypothesized that cells may regulate the intracellular alanine pool by increasing uptake of alanine from the medium. Alanine uptake is mediated by the solute carrier (SLC) family of transporters, with previous studies implicating SNAT1 (SLC38A1), SNAT2 (SLC38A2), ASCT1 (SLC1A4), and ASCT2 (SLC1A5), though SLC1A5 has been reported to have minimal impact on alanine transport 32–34. Compared to other transporters, single-cell transcriptomic data reveal that SLC38A2 is highly expressed in stromal cells, particularly in fibroblasts and myofibroblasts (**Fig. 4A, B**). Despite previous findings indicating that SLC38A2 mRNA levels are not altered in NHLFs treated with TGF-β for 48 h 35, our results demonstrate that TGF-β significantly upregulates SLC38A2 protein expression (**Fig. 4C, D**). Unlike GPT2, the TGF-β-induced upregulation of SLC38A2 protein expression was independent of extracellular glutamine (**Fig. 4C, D**). Previous studies have shown that SLC38A2 protein expression is induced in response to amino acid deprivation 36. We further discovered that supplementation of DMEM with alanine decreased SLC38A2 protein expression in control and TGF-β-treated cells (**Fig. 4E, F**). GPT2 knockdown, which decreases the concentration of intracellular alanine, increased SLC38A2 modestly (**Fig. 4G, H**), suggesting a compensatory upregulation of alanine transport in response to impaired intracellular alanine synthesis.

**Figure 4.**
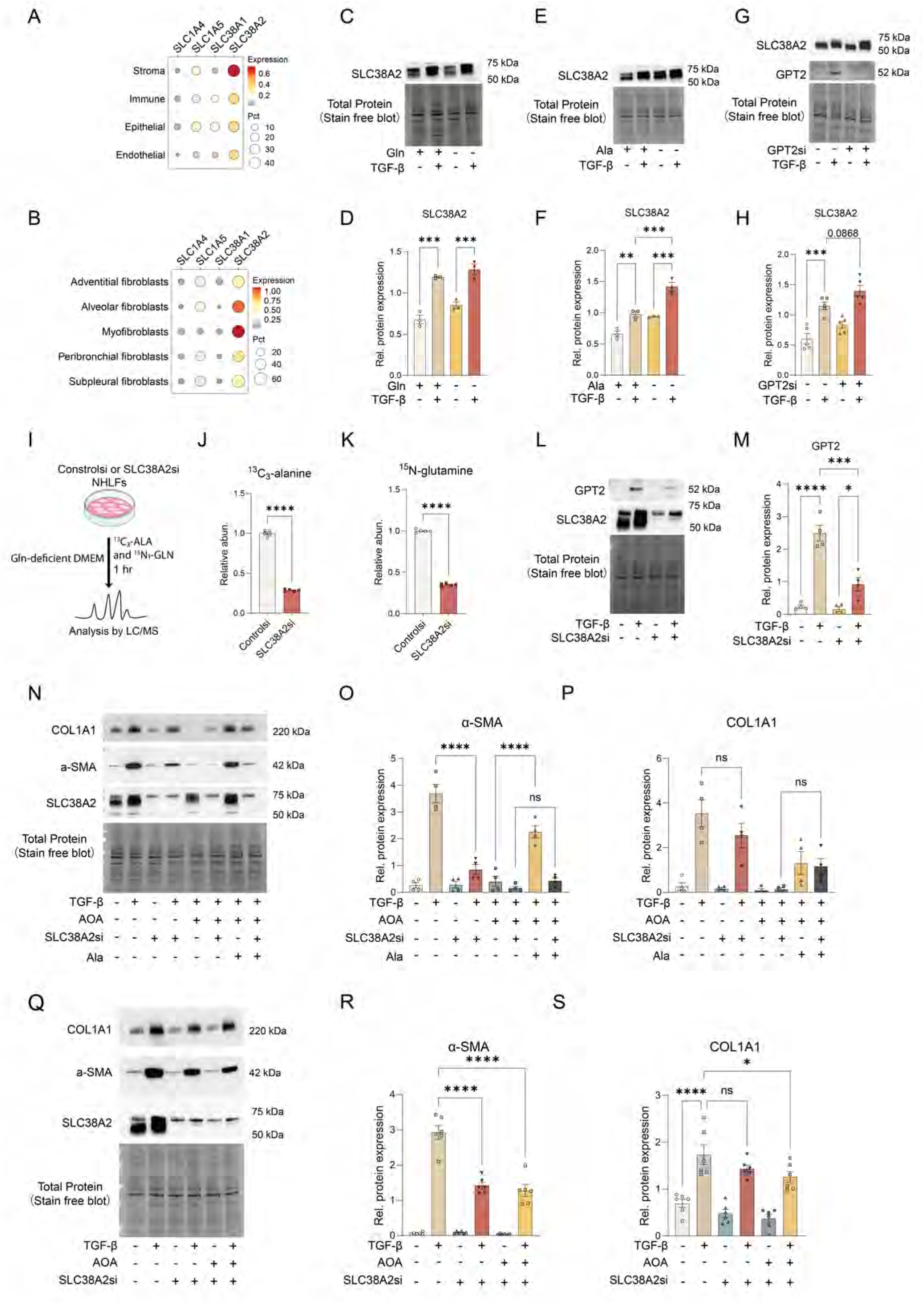
SLC38A2 mediates alanine uptake and myofibroblast differentiation. (A) Normalized transporter expression in lung cell types from the Human Lung Cell Atlas. (B) Fibroblast subset-specific expression in healthy lung samples. Dot color shows mean expression; size shows percent of expressing cells. (C, D) Western blot analysis (C) and quantification (D) showing that TGF-β significantly upregulates SLC38A2 protein expression in NHLF cells, independent of glutamine availability. (E, F) Western blot analysis (E) and quantification (F) showing that extracellular alanine deprivation promotes SLC38A2 protein expression in DMEM-cultured fibroblasts. (G, H) Western blot analysis (G) and quantification (H) showing that GPT2 knockdown reduces intracellular alanine levels and trends toward increasing SLC38A2 protein expression. (I) Schematic overview of the short-term [U-^13^C_3_]-alanine and [^15^N_1_]-glutamine co-uptake assay in control and SLC38A2-knockdown fibroblasts (n = 5 per group). (J, K) Relative quantification of ^13^C_3_-alanine (J) and [^15^N_1_]-glutamine (K) uptake. (L, M) Western blot analysis (L) and quantification (M) showing that SLC38A2 knockdown reduces TGF-β-induced GPT2 expression in FBM. (N-P) Western blot analysis (N) and quantification of α-SMA (O) and COL1A1 (P) expression in control and SLC38A2-knockdown fibroblasts cultured in DMEM for 48 h with TGF-β stimulation, with or without AOA (1 mM) and/or alanine supplementation (2 mM). (Q–S) Western blot analysis (Q) and quantification of α-SMA (R) and COL1A1 (S) expression in FBM-cultured fibroblasts following TGF-β stimulation and combined treatment with AOA and SLC38A2 knockdown for 48 h. For Western blot analysis, individual data points represent biological replicates. J, K: unpaired t-test; D, F, H, M, O, P, R, and S: one-way ANOVA. Data are presented as mean ± SEM. ns, p > 0.05; *p < 0.05; **p < 0.01; ***p < 0.001; ****p < 0.0001.

To determine the relative contribution of SLC38A2 to alanine uptake, we performed short-term uptake assays using [U-^13^C_3_]-alanine in control and SLC38A2-knockdown fibroblasts (**Fig. 4I**). SLC38A2 knockdown decreased alanine import by ∼3-fold, confirming that the majority of alanine uptake is mediated by SLC38A2 (**Fig. 4J**). Interestingly, glutamine uptake was also suppressed by ∼50%, confirming the previously reported role of SLC38A2 as a glutamine transporter 37 (**Fig. 4K**). In line with these findings, SLC38A2 knockdown reduced intracellular alanine and glutamine levels by ∼3- and 2-fold, respectively, after 48 hours of culture in FBM medium (**Fig. EV4A-C**). Notably, glutamine import decreased by 30% when cells were supplemented with 2 mM alanine (**Fig. EV4D, E**), consistent with prior reports of alanine-mediated competitive inhibition of glutamine uptake 31. Given that SLC38A2 functions both as an alanine and glutamine transporter, and that glutamine regulates GPT2 protein expression (**Fig. 2B, C**), we hypothesized that SLC38A2 knockdown would further suppress GPT2 expression and thereby reduce alanine synthesis. As hypothesized, SLC38A2 knockdown significantly reduced the TGF-β-induced GPT2 upregulation in FBM-cultured cells (**Fig. 4L, M**).

We next explored the role of SLC38A2 in myofibroblast differentiation. In DMEM, SLC38A2 knockdown decreased TGF-β-induced upregulation of α-SMA protein expression (**Fig. 4N, O**), possibly by limiting glutamine availability and thereby reducing GPT2-dependent alanine synthesis. This inhibitory effect of SLC38A2 knockdown was potentiated when alanine synthesis was fully blocked by AOA (**Fig. 4N, O**). Notably, alanine supplementation restored α-SMA protein expression in control cells, but not in SLC38A2-knockdown cells, confirming that SLC38A2 is required for alanine uptake-mediated α-SMA protein upregulation (**Fig. 4N, O**). COL1A1 protein expression was only modestly affected by SLC38A2 knockdown (**Fig. 4N, P**). In FBM, SLC38A2 knockdown also decreased the upregulation of α-SMA protein expression after TGF-β treatment, while only modestly decreasing COL1A1 protein expression (**Fig. 4Q-S**), indicating that α-SMA and COL1A1 expression may differ in their reliance on alanine uptake and synthesis. To eliminate intracellular alanine, we combined SLC38A2 knockdown with AOA treatment, which resulted in a significant decrease in COL1A1 protein expression (**Fig. 4Q, S**).

### Alanine deficiency reprograms myofibroblast metabolism

Myofibroblast differentiation in vitro requires increased glycolysis 7,38. To further assess the metabolic consequences of alanine deficiency in myofibroblasts globally, we measured proton efflux rate (PER) and oxygen consumption (OCR) rates following GPT2 knockdown and TGF-β stimulation in DMEM, as indicators of flux through glycolysis and oxidative phosphorylation, respectively. As expected, both PER and OCR were significantly elevated after 48 hours of TGF-β stimulation (**Fig. 5A, B, EV5A-D**). GPT2 knockdown attenuated these effects, reducing both PER and cellular ATP production, primarily through inhibition of glycolysis (**Fig. 5C, D**). Consistently, pharmacological inhibition of GPT2 with BCA and CS produced similar reductions in PER and glycolytic ATP production rates (**Fig. EV5E–J**).

**Figure 5.**
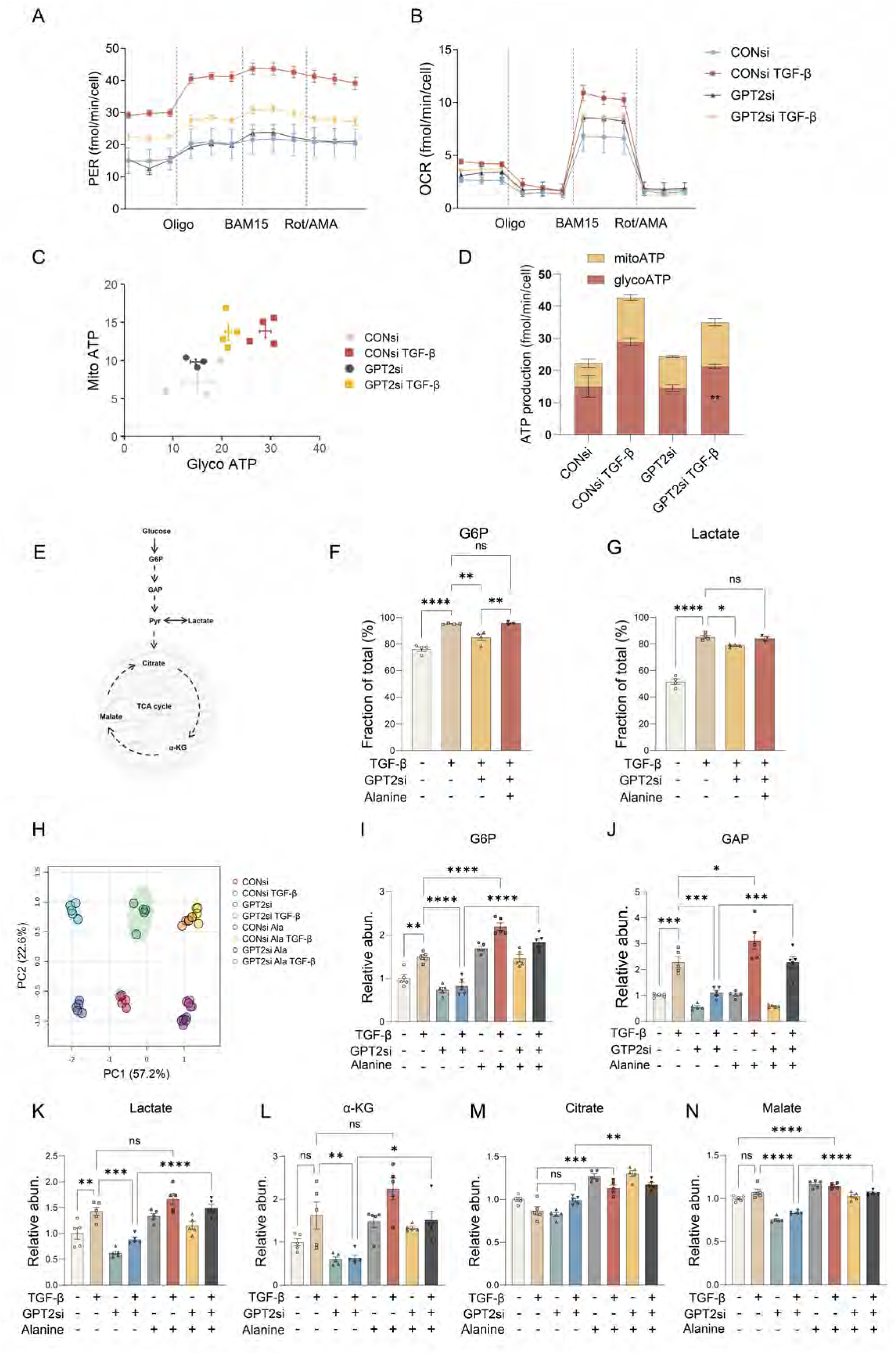
Alanine deficiency reprograms myofibroblast metabolism. (A, B) Proton export rate (PER) and oxygen consumption rate (OCR) in NHLFs treated with TGF-β for 48 h, with or without GPT2 knockdown. (C, D) ATP production from glycolysis and oxidative phosphorylation in NHLFs treated with TGF-β for 48 h, with or without GPT2 knockdown. Data are representative from one of three independent experiments. (E) Schematic overview of glycolysis and the TCA cycle. (F, G) NHLFs cultured in glucose-free DMEM supplemented with glutamine, pyruvate, and 8 mM [U-¹³C₆]-glucose. GPT2 knockdown reduced ¹³C incorporation into G6P (F) and lactate (G), which were restored by alanine supplementation (2 mM). (H) Principal component analysis (PCA) showing metabolic shifts following GPT2 knockdown and alanine supplementation in TGF-β-treated NHLF cells. (I-K) Quantification of intracellular G6P (I), GAP (J), and lactate (K) levels in NHLFs following GPT2 knockdown and alanine supplementation. (L-N) Quantification of intracellular TCA cycle intermediates: α-KG (L), malate (M), and citrate (N). In all cases, control cells were transfected with non-targeting (control) siRNA. D: two-way ANOVA; F, G, I-N: one-way ANOVA with multiple comparisons. A–D show results representative of three independent experiments. Data are presented as mean ± SEM. ns, p > 0.05; *p < 0.05; **p < 0.01; ***p < 0.001; ****p < 0.0001.

To further elucidate the metabolic alterations in glycolysis following TGF-β treatment in the presence or absence of GPT2 knockdown, we conducted metabolic stable-isotope tracing studies (**Fig. 5E**). Lung fibroblasts were cultured in glucose-free DMEM supplemented with [U-^13^C_6_]-glucose (8 mM) and stimulated with TGF-β for 48 h. GPT2 knockdown significantly decreased the labeling enrichment of glucose-6-phosphate (G6P) (**Fig. 5F**) and lactate (**Fig. 5G**) from [U^13^C_6_]-glucose. Alanine supplementation completely or partially restored the enrichment of labeled G6P and lactate, respectively (**Fig. 5F, G**). Metabolomic analysis further confirmed that GPT2 knockdown reshaped the intracellular metabolome, as indicated by the distinct clustering of the different experimental groups in the PCA plot (**Fig. 5H**). Moreover, alanine supplemented control and GPT2 knockdown overlapped, suggesting metabolic rescue (**Fig. 5H**). Specifically, relative to cells transfected with control siRNA, GPT2 knockdown decreased the intracellular abundance of G6P, glyceraldehyde-3-phosphate (GAP), and lactate under TGF-β stimulation, all of which were restored by alanine supplementation (**Fig. 5I-K**). Additionally, GPT2 inhibition decreased the intracellular abundance of several TCA cycle intermediates (*e.g.,* α-KG, malate, and citrate), while alanine supplementation partially rescued their levels (**Fig. 5L-N**). These findings further support an alanine-dependent role for GPT2 in modulating glycolytic and TCA cycle metabolite levels, key metabolic programs previously linked to myofibroblast differentiation 6.

### Alanine metabolism compensates for glutamine deficiency

Myofibroblast differentiation involves both enhanced glycolysis and glutaminolysis ^6^. Glutamine, in particular, contributes to proline biosynthesis, a process required for TGF-β– induced extracellular matrix expression 11,39. Notably, GPT2 knockdown decreased the intracellular concentration of glutamate and proline following TGF-β treatment. These reductions were fully restored upon alanine supplementation (**Fig. EV6A, B**). Given the observation, we hypothesized that alanine may serve as an alternative carbon and nitrogen source for central metabolic intermediates required for myofibroblast differentiation, particularly during partial or complete glutamine deprivation. To test this, we performed stable isotope tracing using [U-^13^C_3_]- or [^15^N]-alanine, in the presence or absence of TGF-β stimulation in DMEM supplemented with or without 2 mM glutamine (**Fig. 6A**). Transamination of [U-^13^C_3_]-alanine gives rise to ^13^C_3_-pyruvate, which can be reduced to ^13^C_3_-lactate by lactate dehydrogenase. Alternatively, ^13^C_3_-pyruvate can be oxidized to ^13^C_2_-acetyl-CoA by pyruvate dehydrogenase or to ^13^C_3_-oxaloacetate by pyruvate carboxylase, yielding ^13^C_2_- and ^13^C_3_-citrate, respectively. Similarly, transamination of [^15^N]-alanine generates [^15^N]-glutamate (**Fig. 6B**). Our data showed that, in the absence of TGF-β, ∼90% of the total intracellular alanine pool was derived from exogenous [U-^13^C_3_]-alanine (**Fig. 6C**). TGF-β treatment decreased the labeling enrichment of alanine to ∼60%, suggesting increased endogenous alanine synthesis from unlabeled substrates during myofibroblast differentiation (**Fig. 6C**). Only a very small fraction of labeled lactate (∼ 2%) was detected (**Fig. 6D**), in line with previous findings that show alanine to be a minor contributor to the total intracellular lactate pool 40. Instead, citrate showed substantial ^13^C-label enrichment (∼15% and ∼40% in the absence or presence of TGF-β, respectively) (**Fig. 6E, EV7A**), suggesting that a greater fraction of the total citrate pool, when compared to the total lactate pool, is derived from alanine. Interestingly, under glutamine-deficient conditions, TGF-β stimulation increased in the labeling of glutamate (∼10% to ∼40%), proline (∼10% to ∼25%), glutamine (∼10% to ∼25%), malate (∼5% to ∼10%), and aspartate (∼10% to ∼20%), indicating that alanine is an important carbon source for these metabolites (**Fig. 6F, G, EV7B-D**).

**Figure 6.**
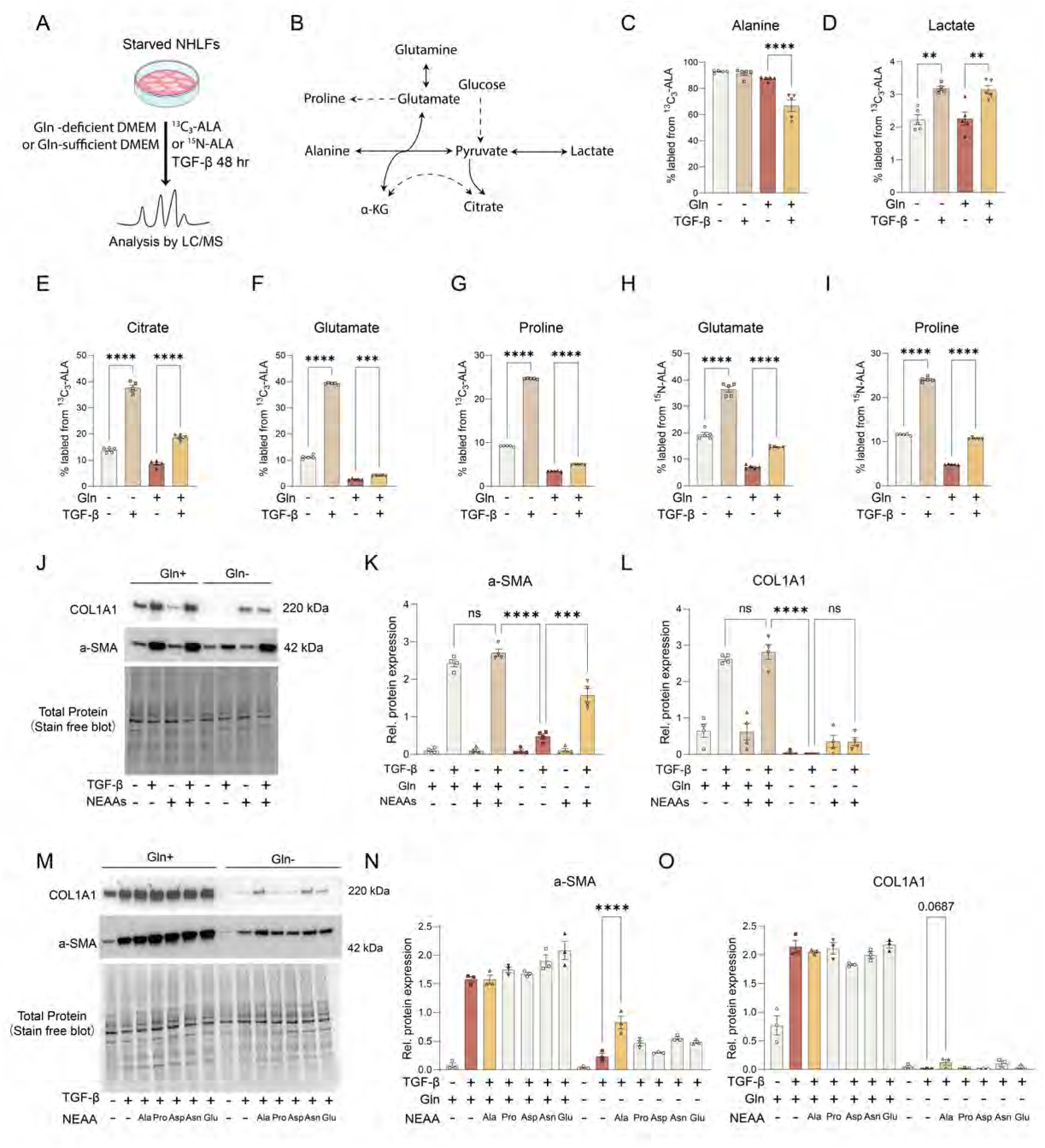
Alanine sustains myofibroblast differentiation by compensating for glutamine deficiency. (A) Schematic overview of isotope tracing experiments using [U-^13^C_3_]-alanine or [^15^N]-alanine in NHLF cells treated with TGF-β in DMEM with or without 2 mM glutamine (n = 5 per group). (B) Expected metabolic fate of alanine. (C) Fractional label enrichment of intracellular ^13^C_3_-alanine derived from extracellular [U-^13^C_3_]-alanine under baseline conditions and following TGF-β stimulation. (D-G) Fractional label enrichment of lactate (D), citrate (E), glutamate (F), and proline (G)from [U-^13^C_3_]-alanine in NHLF cells following TGF-β stimulation for 48 h. (H, I) Fractional label enrichment of ^15^N-glutamate (H) and ^15^N1-proline (I) from ^15^N-alanine under glutamine-sufficient and glutamine-deficient conditions following TGF-β treatment for 48 h. (J) Western blot analysis of α-SMA and COL1A1 expression in fibroblasts cultured in DMEM with or without glutamine following supplementation with a non-essential amino acid (NEAA) cocktail for 48 h. (K, L) Quantification of α-SMA (M) and COL1A1 (N) expression. (M-O) Western blot analysis (M) and quantification of α-SMA (N) and COL1A1(O) expression following individual supplementation of NEAAs (2 mM) in DMEM with or without glutamine. For Western blot analysis, individual data points represent biological replicates. C–I, K, L: one-way ANOVA with multiple comparisons; N, O: one-way ANOVA vs. Gln– TGF-β. Data are presented as mean ± SEM. ns, p > 0.05; *p < 0.05; **p < 0.01; ***p < 0.001; ****p < 0.0001.

In the [^15^N]-alanine experiment, ∼10% of the total glutamate and proline pools were ^15^N-labeled under glutamine-sufficient conditions following TGF-β stimulation, while ^15^N labeling enrichment increased substantially during glutamine deprivation (∼35% for glutamine and glutamate, and ∼25% for proline) (**Fig. 6H, I, EV7E**), demonstrating reverse GPT2 flux allowing transamination of α-ketoglutarate for glutamate and glutamine production.

Since the enhanced synthesis of proline from glutamine facilitates TGF-β-induced matrix protein production, we next investigated whether alanine could compensate during glutamine deficiency to maintain α-SMA and COL1A1 expression. We first applied a commercial NEAA cocktail, supplementation with which was sufficient to, at least in part, rescue the AOA-induced inhibition of TGF-β-induced upregulation of α-SMA and COL1A1 protein expression (**Table 2**, **Fig. EV7F-H**). Under glutamine-deficient conditions, the upregulation of α-SMA and COL1A1 protein expressions following TGF-β stimulation was decreased (**Fig. 6J**); supplementation with NEAAs partially restored α-SMA protein expression, while only modestly effecting COL1A1 production (**Fig. 6J–L**). Next, we supplemented each NEAA individually. Only alanine partially restored the α-SMA protein expression, with minimal change in the COL1A1 protein (**Fig. 6M-O**). These data identify a unique and critical role for alanine as a carbon and nitrogen source for central metabolic intermediates, which may support myofibroblast differentiation under glutamine-deprived conditions.

**Table2.**
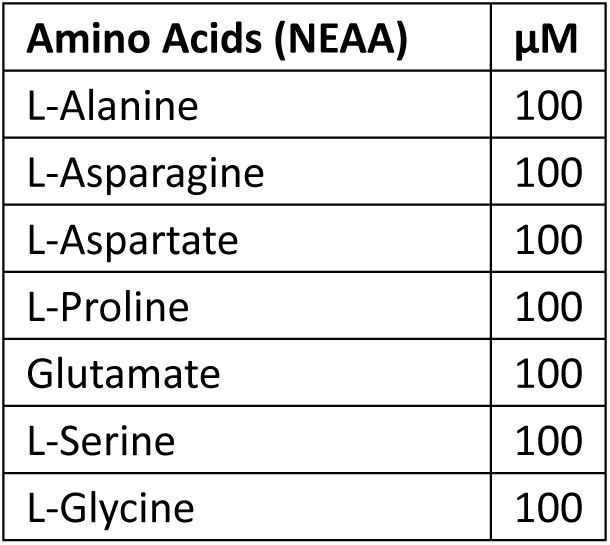
NEAA supplement composition.

| **Amino Acids (NEAA)** | **μM** |
| --- | --- |
| L-Alanine | 100 |
| L-Asparagine | 100 |
| L-Aspartate | 100 |
| L-Proline | 100 |
| Glutamate | 100 |
| L-Serine | 100 |
| L-Glycine | 100 |

### SLC38A2 facilitates myofibroblast function and cooperates with GPT2 to drive TGF-β–induced fibrogenic protein expression

Based on the cooperative regulation of alanine metabolism by GPT2 and SLC38A2 and their essential roles in promoting myofibroblast differentiation, we assessed their clinical relevance in IPF, examined key myofibroblast functional phenotypes, and explored their potential as therapeutic targets. In a publicly available whole-lung RNA-seq dataset (GSE32537), we examined the relationship between SLC38A2 and GPT2 expression and lung function 41. Increased SLC38A2 expression but not GPT2 correlated with more impaired lung function as measured by forced vital capacity (FVC) (**Fig. 7A, EV8**).

**Figure 7.**
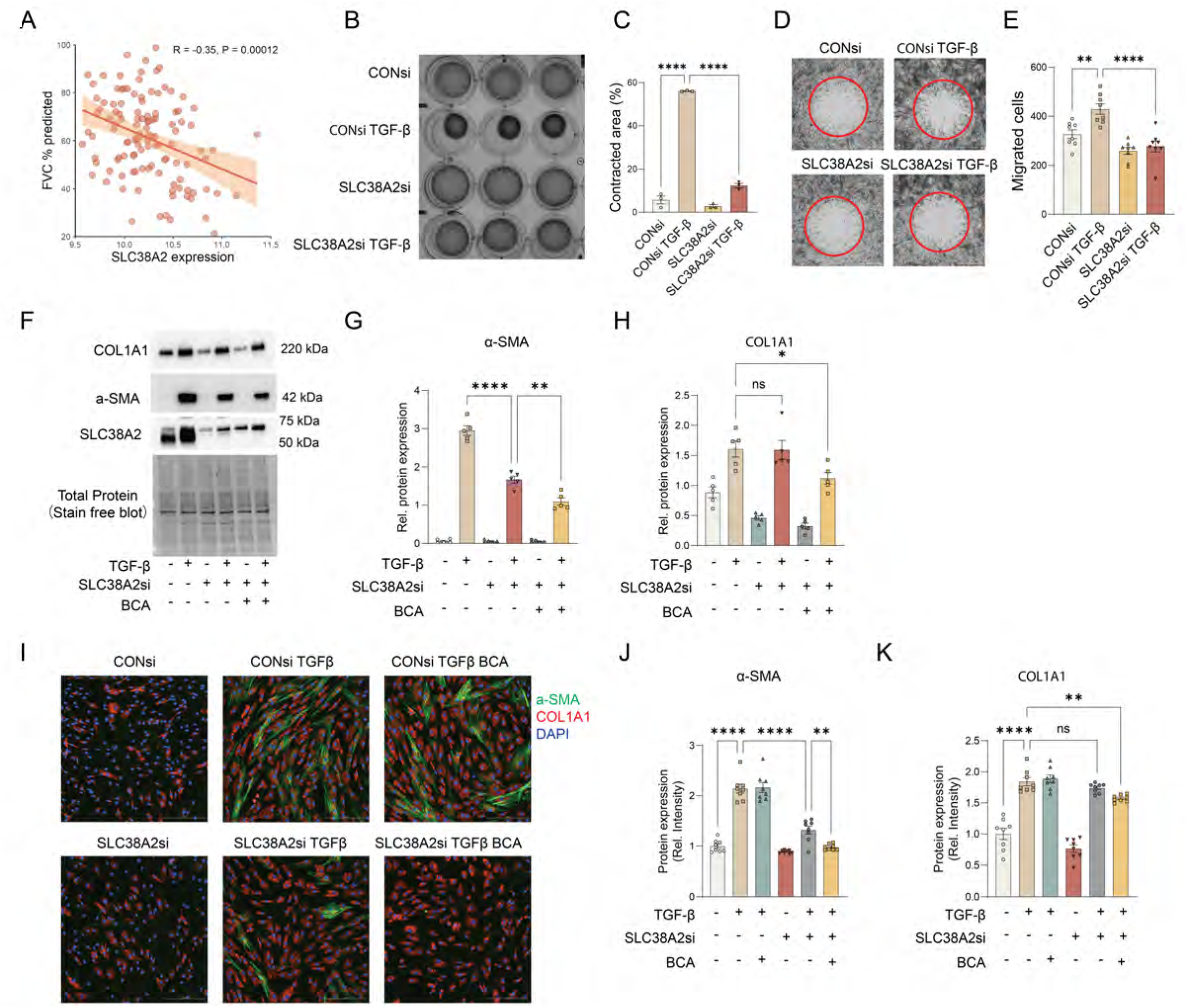
SLC38A2 and GPT2 cooperatively promote TGF-β–induced fibrogenic responses. (A) Pearson’s correlation of SLC38A2 mRNA level and forced vital capacity (FVC) predicted for each patient, from clinical data of GSE32537. (B) Representative images of gel contraction after 24-h TGF-β treatment with or without SLC38A2 knockdown in FBM. (C) Quantification of gel area contraction expressed as the percentage reduction from baseline (0 h) in B. (D) Representative images of control or SLC38A2 knockdown NHLF cells migration into the cell-free zone 24 h after TGF-β treatment in FBM. (E) Quantification of migrated cells within the defined zone based on DAPI staining. (F-H) Western blot analysis (F) and quantification of α-SMA (G) and COL1A1 (H) expression in NHLF cells treated with TGF-β for 48 h, with SLC38A2 knockdown and β-chloro-L-alanine (BCA, 100 µM) treatment. (I-K) Immunofluorescence staining (I) and quantification of α-SMA (J) and COL1A1 (K) expression in IPF patient-derived fibroblasts following SLC38A2 knockdown and BCA (100 µM) treatment in the presence of TGF-β stimulation for 48 h (n = 8 per group). For Western blot analysis, individual data points represent biological replicates. B–E show results representative of three independent experiments. A: linear regression and Pearson correlation. C, E, G, H, J, and K: One-way ANOVA. Data are presented as mean ± SEM. ns, p > 0.05; *p < 0.05; **p < 0.01; ****p < 0.0001.

Myofibroblast differentiation is characterized by enhanced contractile capacity, with α-SMA playing a key role in generating maximal contractile activity 5, 42. Given that SLC38A2 knockdown concurrently impairs both glutamine and alanine uptake and reduces α-SMA expression during myofibroblast differentiation (**Fig. 4Q, R**), we hypothesized that SLC38A2 inhibition may attenuate myofibroblast contractile function. To test this, we evaluated SLC38A2 function in TGF-β–treated fibroblasts cultured in FBM. Notably, SLC38A2 silencing significantly reduced TGF-β–induced gel contraction to near baseline levels (**Fig. 7B, C**). We also assessed fibroblast migratory capacity under the same conditions and found that SLC38A2 knockdown markedly decreased migration, with levels comparable to those of unstimulated controls (**Fig. 7D, E**). Together, these results indicate that SLC38A2 is functionally important for sustaining myofibroblast contractility and motility.

Given that SLC38A2 knockdown only partially reduced GPT2 protein expression, it failed to fully block alanine synthesis (**Fig. 4L, M**). We further explored the feasibility of targeting both alanine synthesis and uptake simultaneously to ameliorate pulmonary fibrosis. In NHLFs, we used SLC38A2 knockdown combined with BCA. Our results show that SLC38A2 knockdown suppressed the TGF-β-induced α-SMA protein expression, and that this suppression was potentiated in the presence of BCA-mediated GPT inhibition (**Fig. 7F, G**). Notably, combined SLC38A2 knockdown and BCA treatment also significantly reduced COL1A1 protein expression (**Fig. 7F, H**). To validate these findings in a disease-relevant context, we performed immunofluorescence staining in fibroblasts derived from IPF patients. The data showed a similar inhibitory effect on α-SMA and COL1A1 protein expression of SLC38A2 knockdown combined with BCA treatment (**Fig. 7I-K**). These findings collectively suggest that dual inhibition of alanine synthesis and transport significantly attenuates TGF-β–induced fibrogenic protein expression.

### Combined inhibition of alanine synthesis and uptake attenuates pulmonary fibrosis

To extend our findings to a more physiologically relevant model, we employed precision-cut lung slices (PCLS), which preserve the native lung architecture and multicellular microenvironment, including both airway and alveolar compartments, making them highly valuable for studying fibrotic pathogenesis, pharmacological responses, and therapeutic discovery in pulmonary fibrosis 43–45. As illustrated in **Fig. 8A**, commercially available PCLS from non-diseased human donors were cultured, transfected with SLC38A2 siRNA, and treated with TGFβ and BCA in DMEM/F-12 medium, which contains both alanine and glutamine (**Table 3**).

**Figure 8.**
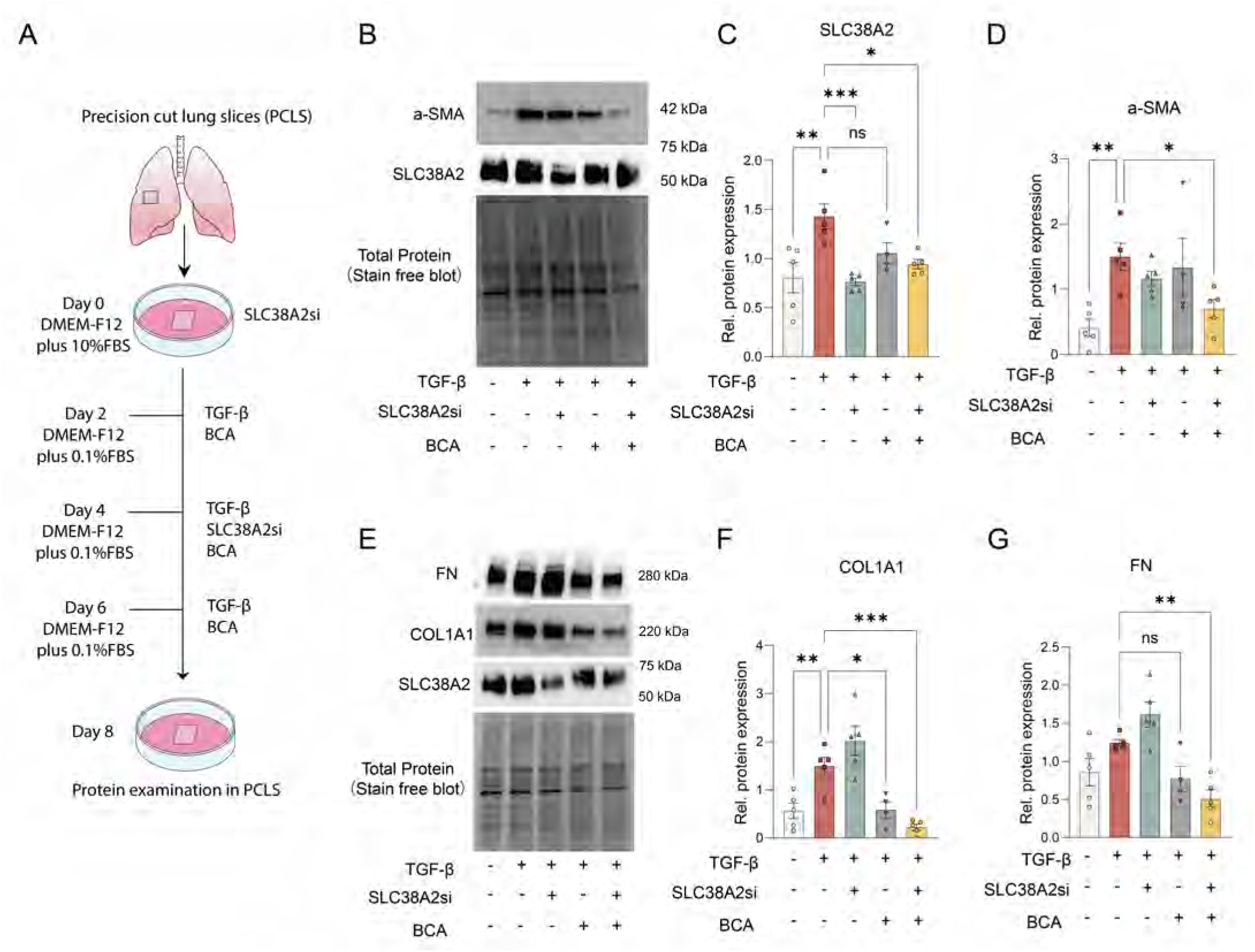
Dual inhibition of alanine uptake and synthesis alleviates pulmonary fibrosis. (A) Schematic overview of experiments using commercially available precision-cut lung slices (PCLS) derived from healthy donors to investigate the role of GPT inhibition and SLC38A2 knockdown in pulmonary fibrosis. In all cases, control PCLS were transfected with non-targeting (control) siRNA. (B-D) Western blot analysis (B) and quantification of SLC38A2 (C) and α-SMA (D) expression in PCLS following TGF-β stimulation with SLC38A2 knockdown and BCA (250 µM) treatment. (E-G) Western blot analysis (E) and quantification of COL1A1 (F) and fibronectin (FN) (G) expression in PCLS following combined SLC38A2 knockdown and BCA (250) µM treatment. For Western blot analysis, individual data points represent biological replicates. C, D, F, and G: One-way ANOVA vs. TGF-β. Data are presented as mean ± SEM. ns, p > 0.05; *p < 0.05; **p < 0.01; ***p < 0.001.

**Table3.** Amino acid composition of DMEM-F12.

| **Amino Acids (DMEM-F12)** | **mM** |
| --- | --- |
| L-Alanine | 0.04999997 |
| L-Arginine hydrochloride | 0.69905216 |
| L-Arginine hydrochloride | 0.05 |
| L-Aspartic acid | 0.05 |
| L-Cysteine hydrochloride-H<sub>2</sub>O | 0.20127796 |
| L-Cystine 2HCl | 0.09996805 |
| L-Glutamic Acid | 0.05 |
| L-Glutamine | 2.5 |
| Glycine | 0.25 |
| L-Histidine hydrochloride-H<sub>2</sub>O | 0.14990476 |
| L-Isoleucine | 0.41580153 |
| L-Leucine | 0.45076334 |
| L-Lysine hydrochloride | 0.4986339 |
| L-Methionine | 0.11570469 |
| L-Phenylalanine | 0.2150303 |
| L-Proline | 0.15 |
| L-Serine | 0.25 |
| L-Threonine | 0.44915968 |
| L-Tryptophan | 0.04421569 |
| L-Tyrosine disodium salt dihydrate | 0.21375479 |
| L-Valine | 0.4517094 |

Consistent with our in vitro findings, SLC38A2 protein expression was upregulated following TGF-β treatment in PCLS, which was effectively suppressed by SLC38A2 knockdown (**Fig. 8B, C**). TGF-β stimulation also induced expression of key fibrogenic proteins in PCLS, including α-SMA, COL1A1 and fibronectin (FN) (**Fig 8D, E-G**). In combination with BCA, SLC38A2 knockdown significantly decreased α-SMA protein levels (**Fig. 8B, D**). Notably, BCA alone decreased the upregulation of COL1A1 following TGF-β treatment; the combined intervention potentiated the decrease in both COL1A1 and FN protein expression (**Fig. 8E-G**). Collectively, these results demonstrate that combined inhibition of alanine synthesis and uptake effectively ameliorates pulmonary fibrosis following TGF-β treatment *ex vivo*.

## Discussion

This study highlights a key role for NEAAs, with particular emphasis on the dynamic regulation of alanine synthesis and uptake, in myofibroblast differentiation. We show that combined intervention strategies targeting alanine synthesis and transport via GPT2 and SLC38A2, respectively, can effectively mitigate the pathological features of myofibroblast differentiation and pulmonary fibrosis.

Myofibroblast differentiation is characterized by increased expression of α-SMA and collagen, accompanied by functional reprogramming including enhanced contractility and migratory capacity. The process relies on key nutrients, particularly glucose and glutamine. Inhibition of glycolysis, or glutamine deprivation, effectively suppress myofibroblast differentiation 16. In this study, we identified alanine as a critical NEAA in myofibroblast differentiation, as other NEAAs did not confer a similar role. Commonly used culture media, including DMEM, do not contain certain amino acids (e.g., alanine), the absence of which affects cellular metabolism and function. For example, the transaminase inhibitor, AOA, suppresses the protein expression of α-SMA in myofibroblasts 10. However, our findings demonstrate that the inhibitory function of AOA on TGF-β-induced upregulation of α-SMA protein expression is attenuated significantly under alanine-replete conditions (*i.e.,* when cells are cultured in FBM or alanine-supplemented DMEM). Medium composition, therefore, is a critical experimental factor in metabolic studies of myofibroblast differentiation. Accurate modeling of metabolic regulation during myofibroblast differentiation in vitro requires the deliberate supplementation, or removal, of specific metabolites (*e.g.,* alanine) to assess their individual contributions.

In addition to glutamine, several other amino acids (*e.g.,* arginine, proline, serine, and glycine) have been reported to play key roles in the progression of pulmonary fibrosis 8,11,46. For example, increased proline synthesis supports TGF-β-induced matrix protein production 11; the serine-glycine pathway supplies glycine for TGF-β1-induced collagen synthesis 8; and the catabolism of glutamine and serine sustains proline and glycine synthesis, thereby promoting collagen production 47. Here, we demonstrate that alanine synthesis intracellularly is indispensable for fibrosis, particularly under conditions of limited alanine uptake. The glutamine-glutamate-α-KG metabolic pathway supports GPT2 expression, and consequently, facilitates alanine production, revealing GPT2-mediated alanine synthesis—tightly coupled to glutamine metabolism—as a key driver of fibrogenesis. Moreover, GPT2 activity is required for the TGF-β–induced upregulation of α-SMA and COL1A1 protein expression by maintaining intracellular alanine levels. Thus, GPT2-dependent alanine synthesis represents an essential and targetable metabolic vulnerability in pulmonary fibrosis.

Alanine homeostasis is maintained by intracellular synthesis and uptake from (and secretion to) extracellular pools, the regulation of which is interrelated. When intracellular synthesis is inhibited, cells may compensate by increasing the uptake from the extracellular milieu, and vice versa. This interlinked regulation of the intracellular and extracellular pools underpins the dynamic regulation of amino acid transporters and their contribution to tumor resistance 48, highlighting the need to target both the intra- and extra-cellular metabolite sources. Our data identify SLC38A2 as a key alanine transporter, and its additional role in glutamine transport may further contribute to its impact on alanine metabolism. Loss of SLC38A2 reduces alanine uptake and decreases glutamine-dependent GPT2 protein expression and alanine synthesis, making it a promising therapeutic target. A previous study demonstrated that inhibition of SLC1A5 with V-9302 targets glutamine metabolism and extracellular matrix deposition effectively in experimental lung fibrosis ^35^, highlighting a potential avenue for antifibrotic therapy involving the targeting of specific amino acid transporters. Interestingly, in addition to SLC1A5, V-9302 has been reported to inhibit SLC38A2-mediated transport 49,50.

Unlike strategies that target glutamine metabolism alone (*e.g.,* the glutaminase inhibitor, CB-839, or SLC1A5 inhibitors), SLC38A2 inhibition not only decreases alanine and glutamine transport directly but also decreases intracellular alanine synthesis indirectly by inhibiting glutamine-dependent GPT2 protein expression and alanine synthesis. As such, SLC38A2 inhibition targets the alanine pools intra- and extra-cellularly, thereby limiting TGF-β-induced myofibroblast differentiation effectively. Nevertheless, we observed that SLC38A2 knockdown alone decreases the TGF-β-induced COL1A1 production only modestly, and this may be explained by SLC38A2-independent alanine or glutamine uptake by cells.

Mechanistically, our data suggest that depleting alanine intracellularly impairs glycolysis. Previous studies have highlighted the contribution of alanine metabolism in tumor, neuronal, and immune cell function 32,40,51. Moreover, GPT2 has been shown to mediate the regulation of glutamine and glucose metabolism by thyroid hormones 52. Our metabolomic data provide mechanistic insight indicating that alanine deficiency decreases the intracellular pools of glycolytic intermediates, including G6P, GAP, and lactate. Alanine augments the TGF-β-induced upregulation of α-SMA and COL1A1 by promoting the production of these intermediates. Alanine may regulate glycolysis by, for example, altering the transcript or protein expression of glycolytic enzymes, modulating glucose uptake, or binding with target proteins that regulate glycolytic flux. Future work should investigate the mechanism by which alanine and its deficiency modulates glycolytic flux.

Glutamine metabolism has emerged as a promising therapeutic target in cancer treatment ^53^. However, therapeutic approaches that target the intracellular pools of certain metabolites are complicated by compensatory roles played by other metabolites (*e.g.,* NEAAs), particularly as a result of flux through converging metabolic pathways. For this reason, cancer cells can resist fluctuations in nutrient availability, thereby undermining the efficacy of single-target metabolic therapies 53. Glutamine metabolism has also been identified as a potential therapeutic target in pulmonary fibrosis 9,10. Yet, as in the case for cancer cells, non-cancer cells (*e.g.,* fibroblasts) likely also exhibit a metabolic flexibility that confers resistance to such therapies. The compensatory roles of NEAAs should be carefully considered. Indeed, our data demonstrate that, under glutamine-deficient conditions, alanine compensates for glutamine as a nitrogen and carbon source to sustain α-SMA protein expression during TGF-β-induced myofibroblast differentiation. Given this metabolic plasticity, a more efficacious therapeutic strategy may be to target glutamine metabolism as well as other key compensatory pathways (*e.g.,* alanine synthesis or uptake) simultaneously. Notably, our study not only uncovers a critical role for alanine metabolism in pulmonary fibrosis but also provides conceptual insights into alanine-centered metabolic adaptation that may be broadly relevant to other pathological contexts, including cancer.

Publicly available transcriptomic data show that SLC38A2 transcript, unlike other alanine transporters, is expressed highly in fibroblasts and correlates with disease severity in IPF patients. This provides a strong rationale for further preclinical investigation focusing on SLC38A2 as a therapeutic target for fibrotic lung disease. Notably, SLC38A2 inhibition alone markedly reduces the contractile and migratory capacities associated with TGF-β–induced myofibroblast differentiation, highlighting the functional sensitivity and therapeutic vulnerability of these processes to nutrient restriction. The use of PCLS as an emerging research model ex vivo enables genetic manipulation or drug intervention while preserving lung tissue structure and microenvironment. Here, we used PCLS to demonstrate that combined inhibition of SLC38A2- and GPT2-mediated alanine uptake and synthesis, respectively, effectively ameliorates the TGF-β-induced upregulation of α-SMA, COL1A1, and FN protein expressions, highlighting the potential efficacy of this combinatorial therapeutic strategy in pulmonary fibrosis. SLC38A2 knockdown alone exhibited only a modest efficacy with respect to suppressing markers of myofibroblast differentiation (*i.e.,* α-SMA and COL1A1), possibly due to a lower knockdown efficiency in PCLS as compared to a monolayer cell culture (*i.e.,* likely a result of less efficient siRNA delivery). This limitation notwithstanding, PCLS remain a valuable translational model in pulmonary research, as they better recapitulate the native cellular microenvironment. Our experiments using PCLSs provide key insights into the *in vivo* relevance of targeting alanine metabolism in IPF.

Moving forward, several key areas warrant further investigation. First, conditional SLC38A2 or GPT2 knockout animal models may be used to validate the role of alanine in fibrosis *in vivo*. Second, current SLC38A2 inhibitors have limited specificity, selectivity, and efficacy 54,55, warranting the development of improved inhibitors. Third, it is important to ascertain whether combined metabolic intervention strategies (*e.g.,* combined BCA and SLC38A2 inhibitors) exhibit synergistic antifibrotic effects *in vivo*. Importantly, our findings propose a broader concept in which disrupting the metabolic flexibility required for fibrogenic responses may represent an effective therapeutic strategy. Further investigations into how various forms of nutrient stress or pharmacological intervention influence cellular metabolic adaptability will be essential for developing therapies that overcome compensatory metabolic rewiring and improve antifibrotic efficacy.

## Methods

### Chemical Reagents

Recombinant human TGF-β1 (CAT #100-21) was purchased from PeproTech, dissolved in 10 mM citric acid, pH 3.0, filtered, and diluted to 10 μg/mL in PBS with 0.1% BSA prior to aliquoting and storing at −80 °C. β-Chloro-L-Alanine (BCA, CAT # HY-107373) was purchased from MedChemExpress and dissolved in water, L-cycloserine (CS, CAT # C1159) was purchased from Sigma Aldrich and dissolved in water, and Aminooxyacetic acid (AOA, CAT # S4989) was purchased from SelleckChem and dissolved in DMSO. Unless otherwise specified, all reagents were used at the indicated concentrations based on previous studies. [U-^13^C_6_]-glucose (CAT # CLM-1396), [U-^13^C_5_]-glutamine (CAT # CLM-1822-H), [α-^15^N_1_]-glutamine (CAT # NLM-557, NLM-1016), [U-^13^C_3_]-L-alanine (CAT # CLM-2184-H), and [^15^N]-L-alanine (CAT # NLM-454) were purchased from Cambridge Isotope Labs. All siRNA was purchased from Dharmacon; anti-SLC38A2 (CAT # BMP081) was purchased from MBL Life Science; anti-GPT (CAT # 16897-1-AP) and anti-GPT2 (CAT # 16757-1-AP) were purchased from Proteintech; anti-FN1 (CAT # ab2413) and anti-α-SMA (CAT #7817) antibodies were purchased from Abcam; anti-Smad3 (CAT # #9523), anti-p-Smad3 (CAT #9520), anti-COL1A1 (CAT # 39952), HRP-anti-Rabbit IgG (CAT #7074), and HRP-anti-Mouse IgG (CAT #7076) were obtained from Cell Signaling Technology.

### Fibroblast cell culture

Primary normal human lung fibroblasts (NHLFs) were obtained from Lonza (CAT # CC-2512), cryopreserved at passage 3, and used between passages 3 and 8. Cells were cultured in FGM-2 medium, consisting of FBM basal medium (Lonza, CAT# CC-3131) supplemented with FBM SingleQuots™ Supplements and Growth Factors (Lonza, CAT# CC-4126), and subjected to starvation in serum-free media (DMEM or FBM) for 24 hours unless otherwise specified. Cells were then treated with recombinant human TGF-β1 (2 ng/mL) for 48 h to induce myofibroblast differentiation in vitro.

Lung fibroblasts from IPF patients were derived from lung transplantation surgeries. Program and their collection were approved by the Mass General Brigham Institutional Review Board (2013P002332, 2016P001890, 2019P003592, 2020P002765). Lung tissues were minced and digested with Liberase (Sigma, CAT # 05401020001) and DNase I (Sigma, CAT # 10104159001), followed by filtration using a sterile 70 μm filter. The filtrates were centrifuged at 300×g, washed once with RPMI medium (Lonza, CAT# RPMI), and plated in DMEM supplemented with 10% (v/v) FBS, penicillin, and streptomycin. After two passages, the medium was replaced with FGM-2 medium. Cells were cryopreserved in liquid N2 at passage 3 and used for experiments at passage 4.

### Isotope Labeling Experiments

For alanine uptake experiments, cells treated with TGF-β for 24 h were switched to fresh DMEM supplemented with 2 mM ^13^C_3_-L-alanine. For glutamine and alanine co-uptake experiments, cells subjected to SLC38A2 knockdown for 48 h were switched to fresh glutamine-free DMEM containing 2 mM ^13^C_3_-L-alanine and 2 mM [α-^15^N_1_]-glutamine. Similarly, for glutamine uptake experiments, cells pre-treated with 2 mM unlabeled alanine in DMEM for 24 h were switched to fresh glutamine-free DMEM containing 2 mM ^13^C_5_-glutamine glutamine and 2 mM unlabeled alanine. After 1 h of incubation, cells were washed three times with LC-MS-grade water (Thermo Fisher Scientific), and lysates were prepared for metabolomic analysis as described below.

To study alanine synthesis, cells starved in DMEM for 24 h were switched to glucose-free DMEM supplemented with ^13^C_6_-glucose, glutamine-free DMEM supplemented with ^13^C_5_-glutamine, or glutamine-free DMEM supplemented with [α-^15^N_1_]-glutamine. After 48 h, cells were washed with LC-MS-grade water twice, and lysates were prepared for metabolomic analysis as described below.

Alternatively, to study the metabolic fate of alanine, cells starved in DMEM for 24 h were switched to glutamine-free DMEM supplemented with 2 mM ^13^C_3_-L-alanine or ^15^N-L-alanine, with or without 2 mM glutamine, and treated with TGF-β for 48 h. Cells were then washed twice with LC-MS-grade water, and lysates were prepared for metabolomic analysis as described below.

### Western Blotting

Cells were lysed on ice in buffer containing Tris 10 mM, pH 7.4, NaCl 150 mM, EDTA 1 mM, EGTA 1 mM, Triton X-100 1% v/v, NP-40 0.5% v/v, and 1x Halt™ Protease and Phosphatase Inhibitor Cocktail (ThermoFisher Scientific, CAT# 78440). Protein concentrations were measured using the bicinchoninic acid (BCA) assay (ThermoFisher Scientific, CAT# 23227). Lysates were standardized to 10–20 µg of protein (30 µg of protein was used for the p-Smad3 immunoblot), separated by SDS-PAGE on stain-free Tris-glycine gels (Bio-Rad), cross-linked by 45 s of UV illumination, and imaged using the ChemiDoc system (Bio-Rad). Proteins were transferred to PVDF membranes using the Trans-Blot Turbo transfer system (Bio-Rad). Stain-free membrane images were used to assess total protein loading.

Membranes were blocked in 5% (v/v) milk (Bio-Rad, CAT# 170-6404), incubated with primary antibodies overnight at 4 °C, followed by incubation with secondary antibodies at room temperature for 1 h. Bands were visualized using WesternBright ECL reagent (Advansta, CAT# K-12045-D20). Band intensities were quantified using Image Lab software (Bio-Rad) and normalized to total protein signals.

### RNA Interference

GPT1 (CAT# L-031622-01), GPT2 (CAT# L-004173-01), or SLC38A2 (CAT# L-007559-01) siRNA (Dharmacon) was reverse transfected into ∼4 × 10⁵ lung fibroblasts per well in 6-well plates using 20 pmol siRNA and 7.5 µL Lipofectamine RNAiMAX transfection reagent (Thermo Fisher Scientific, CAT# 13778). Non-targeting siRNA was used as a transfection control (CAT# D-001810-10-05). After 24 h of transfection, cells were serum-starved for an additional 24 h, followed by TGF-β treatment (2 ng/mL).

### Immunofluorescence Staining

Cells were seeded at 8,750 cells/well in 96-well black-wall, clear-bottom plates. After 24 h, cells were serum-starved for 24 h and treated with TGF-β and inhibitors for 48 h. For combined SLC38A2 knockdown and GPT inhibition treatment, cells were transfected for 24 h, seeded, and starved for 24 h. After treatments, cells were washed with PBS, fixed in 4% (v/v) paraformaldehyde for 15 min, and permeabilized with 0.5% (v/v) Triton X-100 for an additional 15 min. Blocking was performed with 3% (v/v) bovine serum albumin (BSA, Sigma, CAT# A9576) for 30 min at room temperature. Cells were then incubated with primary antibodies against α-SMA (Abcam, CAT# ab7817, 1:1500) and COL1A1 (Cell Signaling Technology, CAT# 39952, 1:1000) at room temperature for 2 h. After three washes with PBS, the cells were incubated with fluorescently labeled secondary antibodies (goat anti-mouse Alexa Fluor 488 (Thermo Fisher Scientific, CAT# A11001, 1:1000) and goat anti-rabbit Alexa Fluor 647 (Thermo Fisher Scientific, CAT# A21244, 1:1000)) for an additional 45 min at room temperature. After three washes with PBS, nuclei were stained with 4’,6-diamidino-2-phenylindole (DAPI, 2ug/ml) for 10 min, followed by three PBS washes. Immunofluorescence images were acquired using a Cytation™ 5 Cell Imaging Multi-Mode Reader (Agilent BioTek Instruments, USA). Cells were imaged using a 20× objective for representative images and a 4× objective for quantitative analysis, with filter channels for DAPI, Alexa Fluor 488, and Alexa Fluor 647. Exposure time and gain settings were kept consistent across all samples. Image acquisition and analysis including cell counting and total fluorescence intensity per cell were performed automatically using Gen5™ software.

### Seahorse Assay

Lung fibroblasts transfected with GPT2 siRNA for 24 h were seeded at 20,000 cells/well in 24-well Seahorse plates. After 24 h of serum starvation and subsequent 48 h of TGF-β treatment, the medium was replaced with seahorse DMEM (Agilent, CAT# 103335-100) supplemented with 10 mM glucose, 2 mM glutamine, and 1 mM pyruvate. Basal oxygen consumption rate (OCR) and proton export rate (PER) were measured using an XFe24 analyzer (Agilent), with sequential injections of oligomycin (1 µM), BAM15 (2.5 µM), and rotenone/antimycin A (0.5 µM each). After the assay, cells were stained with Nuclear Green LCS1 (10 µM; Abcam CAT# ab138904) for 30 min and counted using a Cytation 5 imager for normalization.

### Metabolomics

#### Sample preparation

Extracellular metabolites were extracted by mixing conditioned media with pre-cooled LC-MS-grade methanol (Honeywell, CAT# 34966,) at a 1:4 (vol:vol) media:methanol ratio on ice. For analysis of intracellular metabolites, cells were rapidly washed twice with HPLC-grade water and snap-frozen by placing the plates on liquid N_2_. Plates were stored at −80 °C until metabolite extraction. Metabolites were extracted with 500 μL of extraction buffer [50:30:20 (vol:vol:vol) LC-MS-grade methanol:acetonitrile:water containing 1 μM D_8_-valine and ^13^C_5_-glutamine as internal standards (internal standards were not added for tracing experiments)] at −80 °C. Samples were centrifuged at 17,000 × *g* for 10 min at 4 °C. Supernatants were dried using a SpeedVac concentrator (Thermo Savant) at 42 °C, reconstituted in 50 μL of HPLC-grade water, centrifuged at 17,000 × *g* for 10 min at 4 °C, and transferred (35 μL) to LC-MS vials.

#### LC-MS data acquisition and analysis

LC-MS analyses were performed using a Vanquish UHPLC system connected to a Q Exactive Orbitrap mass spectrometer with a HESI-II source (Thermo Fisher Scientific). Samples were stored in the autosampler at 5 °C prior to injection. The following chromatographic parameters were used: solvent A, 20 mM ammonium carbonate supplemented with 5 μM medronic acid, pH corrected to 9.2 with ammonium hydroxide; solvent B, acetonitrile; injection volume, 2 μL; oven temperature, 25 °C; flow rate, 0.1 mL/min. Compounds were separated using a ZIC-pHILIC stationary phase (150 mm × 2.1 mm × 3.5 mm; Merck) with a guard column and a linear gradient of solvent B as follows: 0 min, 80%; 20 min, 20%; 20.5 min, 8%; 24 min, 8%; 24.5 min; 80%; 35 min, 80%. MS scan parameters were as follows: scan type, full MS; scan range, 60-900 Th; fragmentation, none; resolution, 70,000; microscans, 1; lock masses, off; AGC target, 1 x 106; maximum injection time, 80 ms. The MS was operated in polarity switching mode. HESI source parameters were as follows: sheath gas flow rate, 40, auxiliary gas flow rate, 15; sweep gas flow rate, 1; spray voltage, 1 kV; capillary temperature, 320 °C; S-lens RF level, 50; auxiliary gas heater temperature, 350 °C. Peak identification was performed using the Thermo TraceFinder General LC software (Thermo Fisher Scientific), using an in-house library of metabolites with known retention times, as assessed and optimized previously with authentic standards using the same method parameters. To correct for total signal variation across samples, areas under the curve (AUCs) of integrated peaks were normalized to the total sum of all AUCs in the same sample. Volcano plots, heatmaps, and KEGG pathway enrichment plots were generated using R (version 4.4.3) with relevant packages.

### Gel contraction assay

NHLF cells were transfected with either control siRNA or SLC38A2 siRNA for 24 h, followed by resuspension in FBM and neutralized collagen solution (TeloCol-3, Advanced Biomatrix) at a 1:2 ratio to yield a final concentration of 100,000 cells/mL and 1.8 mg/mL collagen. The mixture (0.5 mL) was added to 24-well plates and allowed to harden for 10 min at 37 °C, after which 1 mL of FBM was added. Cells were serum-starved overnight prior to gel release. Baseline (0 h) images were acquired using a ChemiDoc imager (Bio-Rad), followed by treatment with TGF-β (2 ng/mL). After 24 h, images were acquired, and gel areas were measured using Image Lab software (Bio-Rad). Data were expressed as the percentage of gel contraction relative to the initial area at 0 h using the formula: [(Areat=0 – Areat=24) / Areat=0] × 100.

### Cell migration assay

NHLF cells were transfected with either control siRNA or SLC38A2 siRNA for 24 h, then seeded at a density of 1 × 10⁵ cells/mL (100 µL per well) into 96-well migration plates equipped with stoppers (Platypus Technologies, CMA1.101). After overnight serum starvation, the stoppers were removed, baseline (0 h) images were acquired using the white light channel of the Cytation™ 5 Cell Imaging Multi-Mode Reader, and the initial cell-free migration zone was defined. Cells were then treated with TGF-β (5 ng/mL) and allowed to migrate for 24 h. After migration, cells were fixed and permeabilized with 4% (v/v) paraformaldehyde containing 0.5% (v/v) Triton X-100 for 15 min, followed by staining with DAPI. Fluorescence images were acquired using the DAPI channel of the Cytation 5 system. The number of cells that migrated into the defined zone was quantified using ImageJ. For representative visualization, selected wells were stained with 0.1% (v/v) crystal violet for 15 min and imaged using the white light channel of the Cytation 5 system.

### Precision-Cut Lung Slices

PCLS were obtained from Anobios and cultured in DMEM/F12 medium containing 10% (v/v) FBS for 48 h. siRNA transfections and TGF-β/inhibitor treatments were performed, with fresh media and treatments replenished every 48 h. On day 6, slices were collected, lysed in radioimmunoprecipitation (RIPA) lysis buffer with protease inhibitors and homogenized using bead beating for protein extraction. Western blotting was performed as described above. Experiments included five biological replicates.

### Statistical Analyses

Unless otherwise stated, data analysis, visualization, and statistical comparisons were performed using GraphPad Prism 10. Quantification of Western blots was based on at least three biological replicates. Seahorse and other functional assays were independently performed in a minimum of three biological replicates, and representative data from a single experiment are shown. Data are presented as mean ± SEM. Normality was assessed using the Shapiro–Wilk test. Two-group comparisons were performed using two-tailed unpaired t-tests, Welch’s t-tests, or Mann–Whitney U tests, as appropriate. For comparisons among multiple groups, one-way ANOVA followed by Tukey’s or Bonferroni post hoc test was used. For experiments involving two independent variables, two-way ANOVA with Tukey’s post hoc test was applied. Correlations were assessed using linear regression and Pearson correlation. P values < 0.05 were considered statistically significant.

## Acknowledgments

This work was supported by project grants from National Institutes of Health grant R01HL167718 (WMO)

## Author contributions

Conceptualization: FL, WMO

Methodology: FL, NV, WMO

Investigation: FL, NV, DRZ, WMO

Visualization: FL, NV, WMO

Funding acquisition: WMO

Project administration: WMO

Resources: DRZ, WMO

Supervision: WMO

Writing - original draft: FL, NV

Writing - review and editing: FL, NV, DRZ, MK, HNN, EYK, MLS, WMO

## Data availability

All data associated with this study are available in the main text or the supplementary materials. Additional information is available from the corresponding author upon reasonable request.

---

**Figure EV1.**
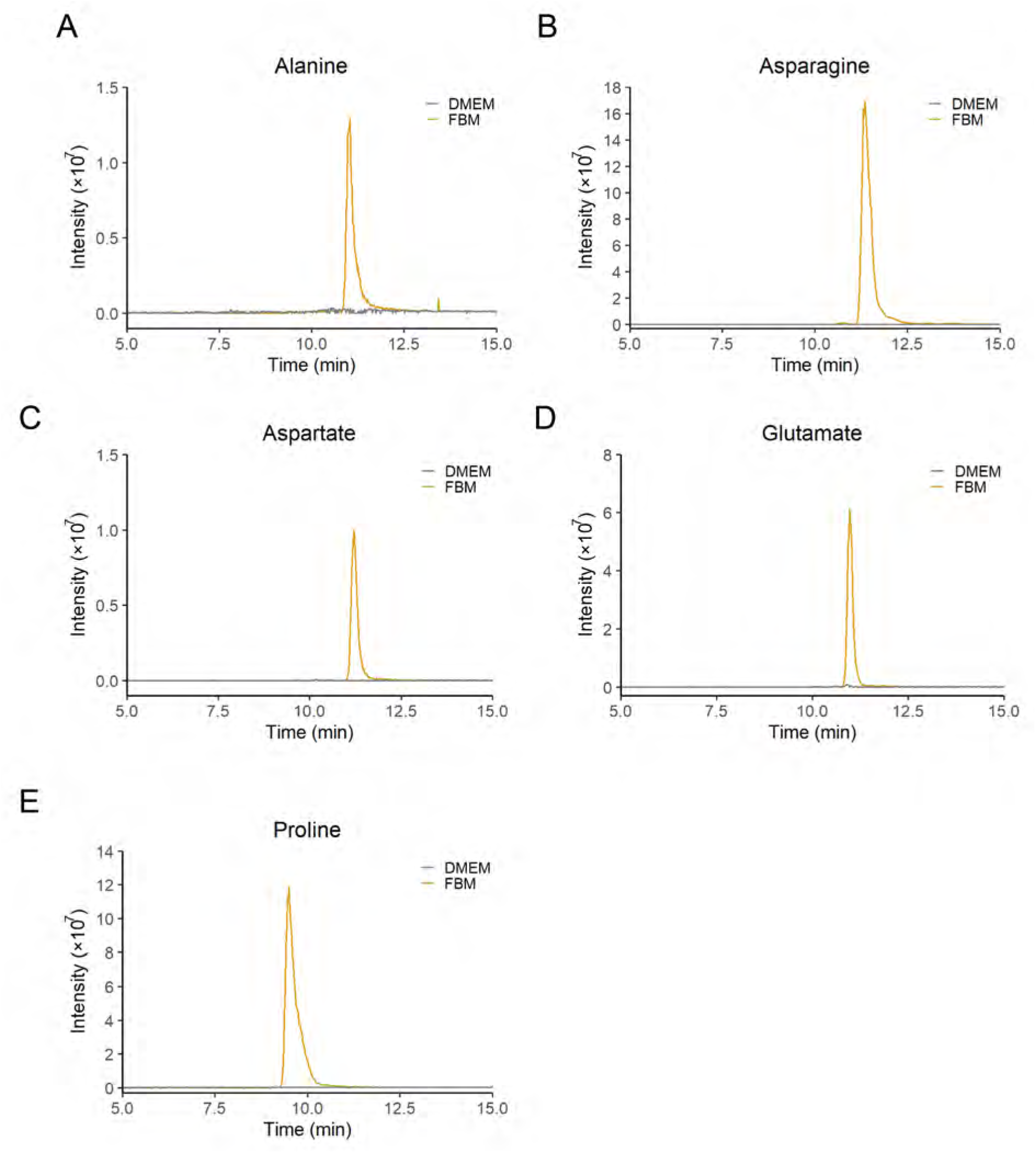
Amino acid composition of DMEM and FBM. (A-E) Representative LC-MS chromatograms of selected NEAAs in FBM and DMEM. Shown are extracted ion chromatograms (± 3 ppm) for alanine (m/z 90.0550, RT 11.0 min), asparagine (m/z 133.0608, RT 11.4 min), aspartate (m/z 134.0448, RT 11.2 min), glutamate (m/z 148.0604, RT 11.0 min), and proline (m/z 116.0706, RT 9.5 min).

**Figure EV2.**
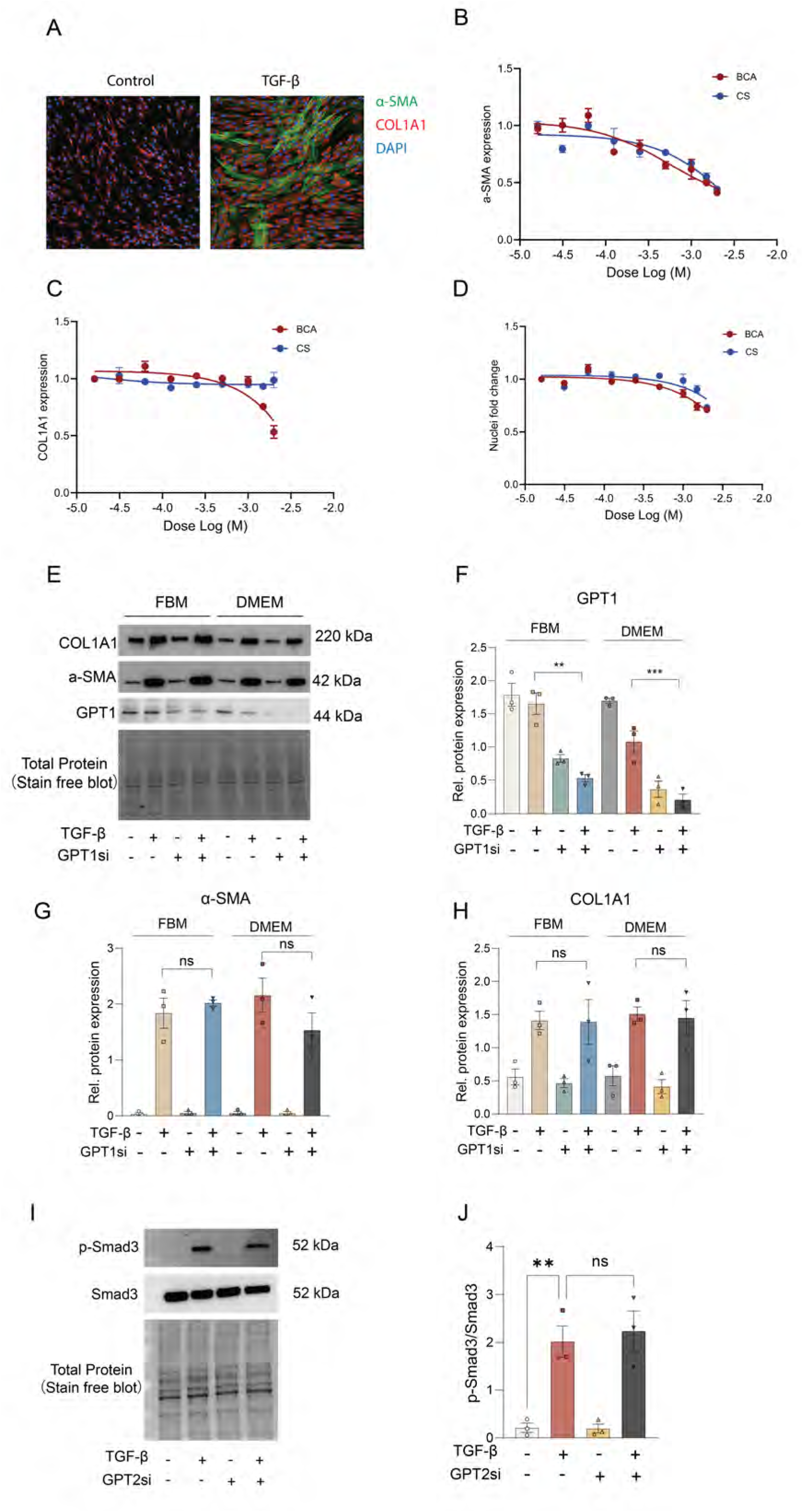
Effects of GPT inhibition on myofibroblast differentiation. (A) Representative immunofluorescence images showing α-SMA (green), COL1A1 (red), and DAPI (blue) in control and TGF-β–treated NHLFs. (B–D) Immunofluorescence quantification showing the dose-dependent effects of GPT1/2 inhibitors L-cycloserine (CS) and β-chloro-L-alanine (BCA) on α-SMA expression (B), COL1A1 expression (C), and cell number (D) as assessed by DAPI-stained nuclei in FBM-cultured NHLFs after 48 h of treatment. (E–H) Western blot analysis showing that GPT1 knockdown does not significantly affect α-SMA or COL1A1 expression in either FBM or DMEM medium. Quantification of GPT1 (F), α-SMA (G), and COL1A1 (H) protein levels is shown. (I, J) Western blot analysis showing that GPT2 knockdown does not alter Smad3 phosphorylation (p-Smad3) after 1 h TGF-β treatment. Quantification of p-Smad3/Smad3 ratio is shown in (J). For Western blot analysis, individual data points represent biological replicates. F-H, and J: One-way ANOVA. Data are presented as mean ± SEM. ns, p > 0.05; **p < 0.01; ***p < 0.001.

**Figure EV3.**
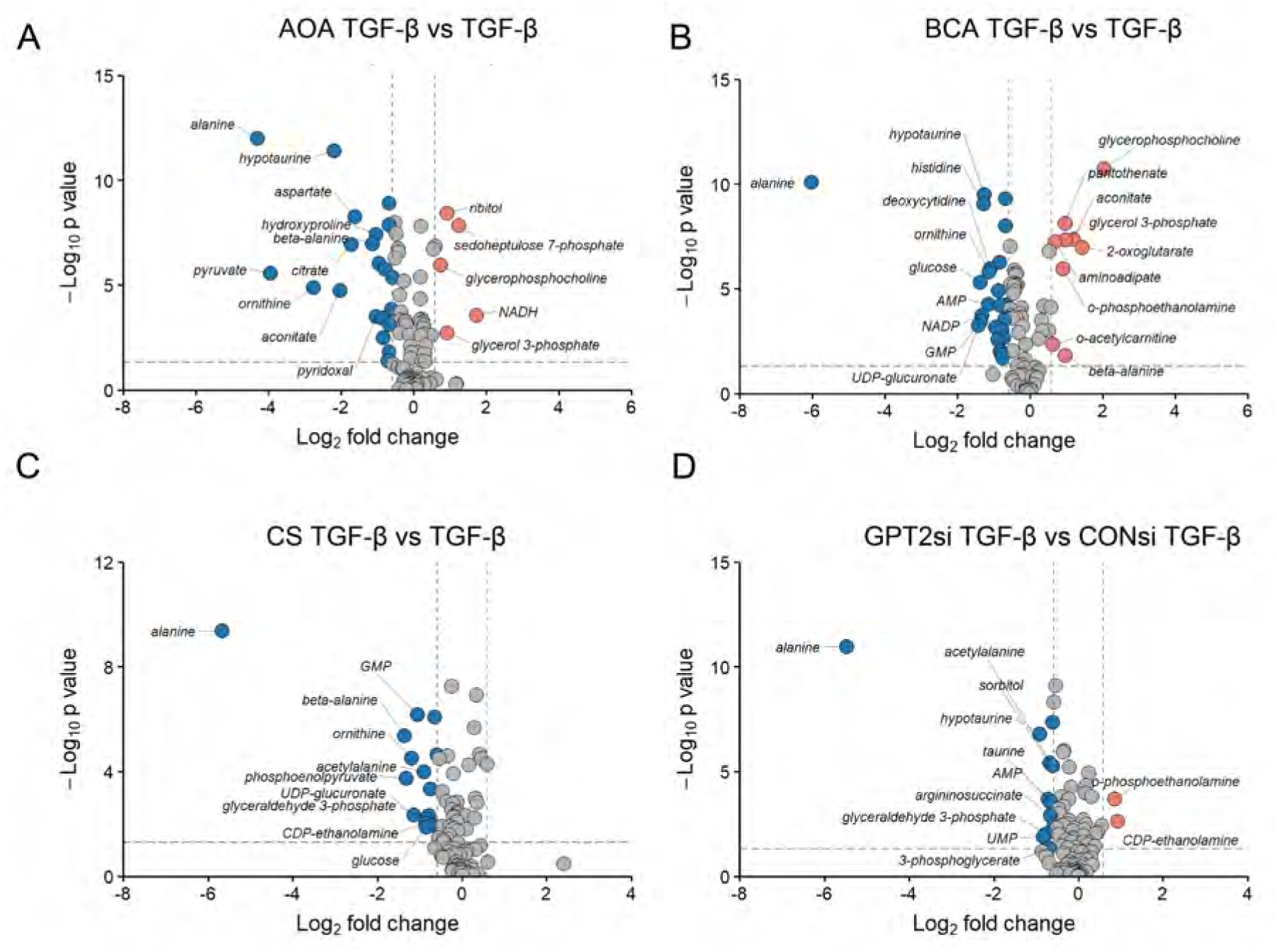
Effects of GPT2 inhibition on profibrotic metabolites. (A–D) Volcano plots showing metabolomic changes in TGF-β–treated NHLF cells upon inhibition or knockdown of GPT2 using AOA (A), BCA (B), CS (C), or GPT2 siRNA (D). Differential metabolites were selected based on a fold change (FC) threshold ≥ 1.5 and a P-value ≤ 0.05. The top 10 most significantly altered metabolites are labeled.

**Figure EV4.**
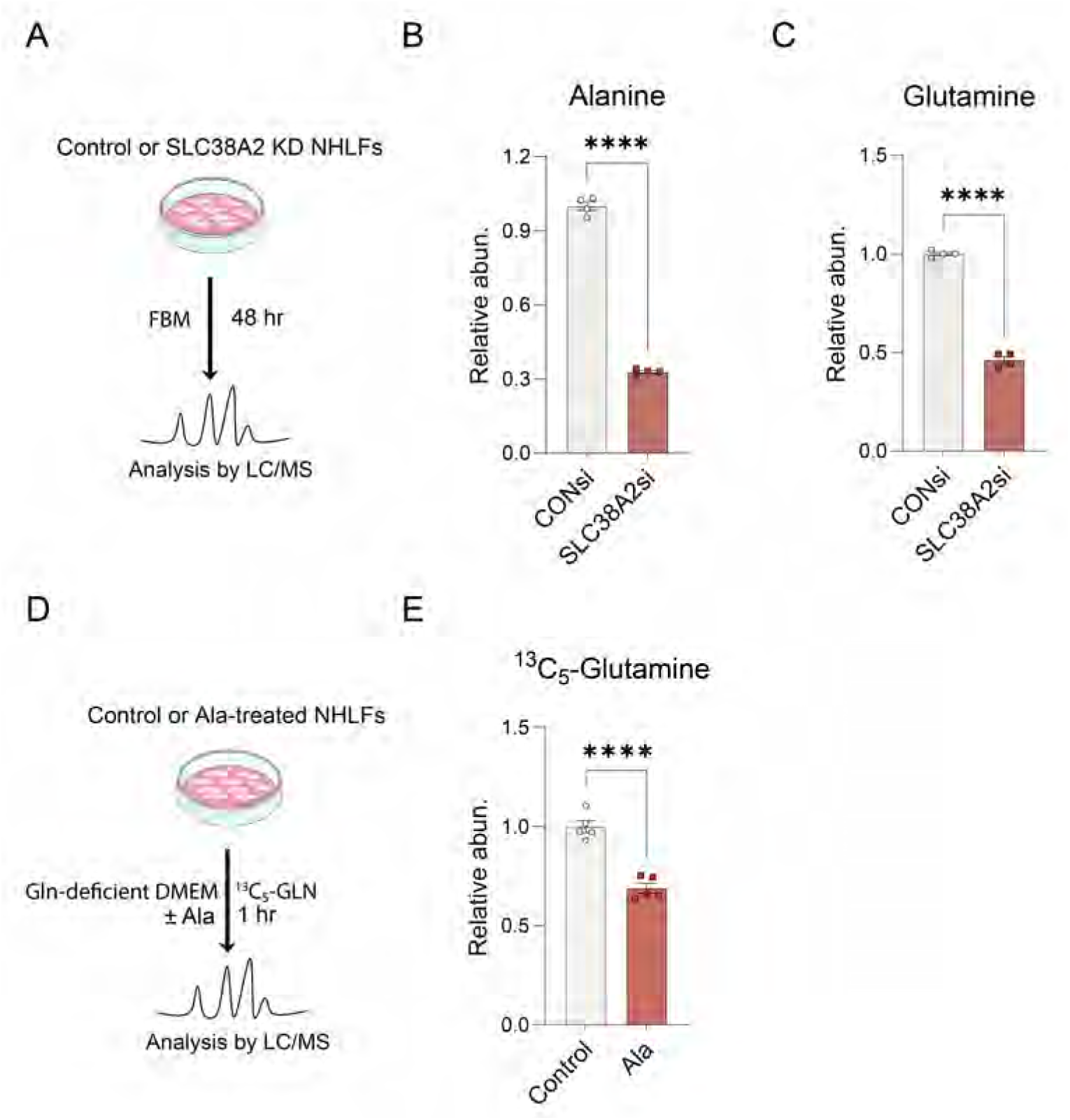
SLC38A2 mediates alanine-supported myofibroblast activation and modulates glutamine uptake. (A) Experimental schematic for measuring intracellular alanine and glutamine in control and SLC38A2-knockdown NHLFs after 48 hours of FBM culture. (B, C) Relative intracellular levels of alanine (B) and glutamine (C) in the indicated conditions. (D) Schematic of the experimental design to assess the effect of extracellular alanine (2 mM) on [^13^C_5_]-glutamine uptake in glutamine-deficient DMEM. (E) Relative quantification of intracellular ^13^C_5_-glutamine levels showing that alanine supplementation significantly reduces glutamine uptake. B, C, and E: unpaired t-test; Data are presented as mean ± SEM. ****p < 0.0001.

**Figure EV5.**
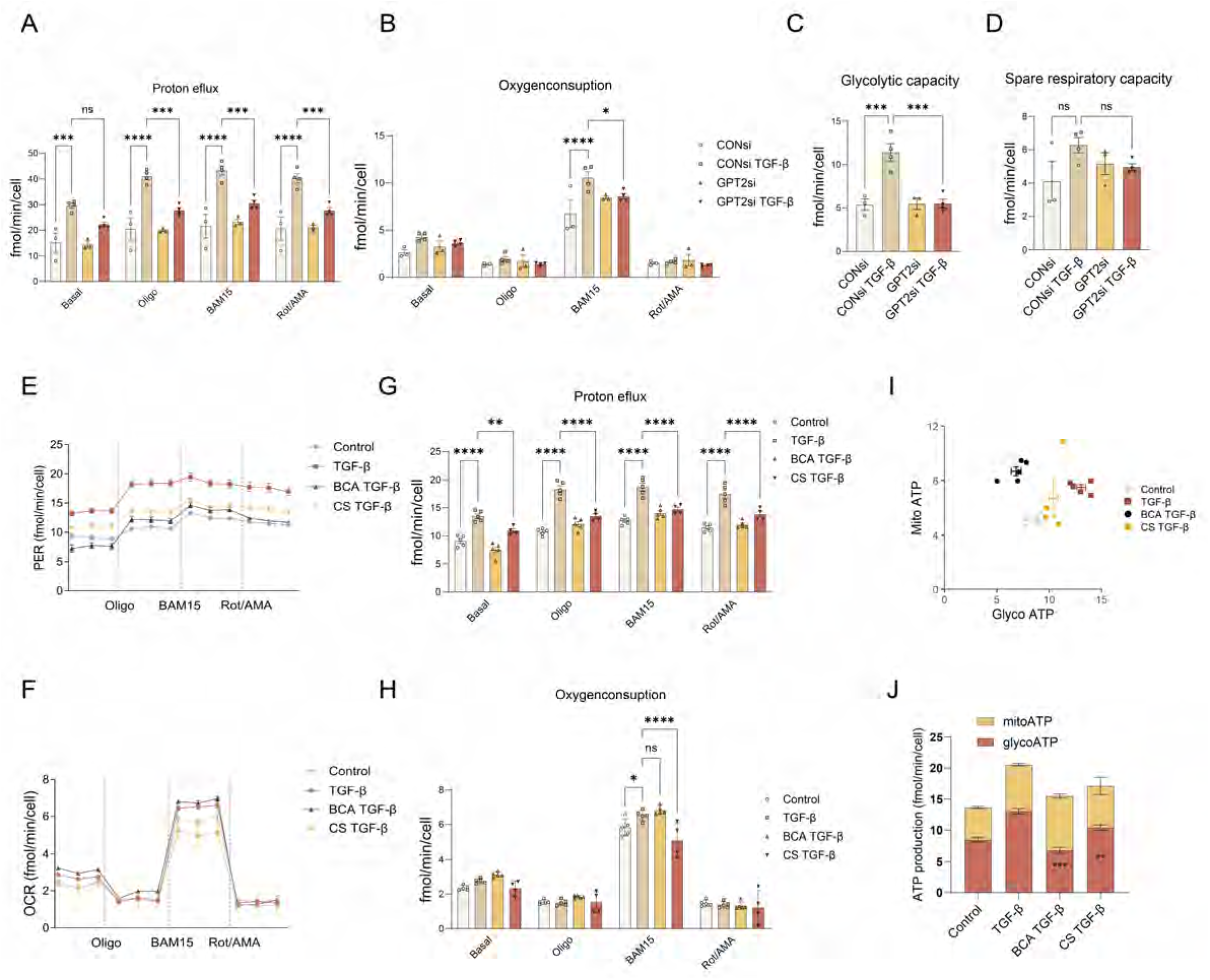
GPT2 inhibition disrupts cellular bioenergetics and anabolic metabolism. (A, B) Summary of proton efflux (PER) and oxygen consumption rates (OCR) from Seahorse XF analysis in NHLF cells with or without GPT2 knockdown and TGF-β treatment. Quantification is based on Fig. 5A and 5B. (C, D) Cell bioenergetic stress test results: glycolytic capacity was determined by the increase in proton efflux rate after oligomycin treatment; spare respiratory capacity was calculated as the increase in OCR after BAM15 treatment relative to basal levels. (E, F) PER (E) and OCR (F) in NHLF cells treated with TGF-β (48 h) in DMEM, with or without pharmacological inhibition of GPT2 using BCA or CS. (G, H) Summary of Seahorse analysis showing PER and OCR across all conditions. (I, J) Glycolytic and mitochondrial ATP production in NHLFs treated with TGF-β (48 h) in DMEM with or without GPT2 inhibition. A, B, G, H, and J: Two-way ANOVA, C and D: One-way ANOVA. Data are presented as mean ± SEM. ns, p > 0.05; *p < 0.05; ***p < 0.001; ****p < 0.0001.

**Figure EV6.**
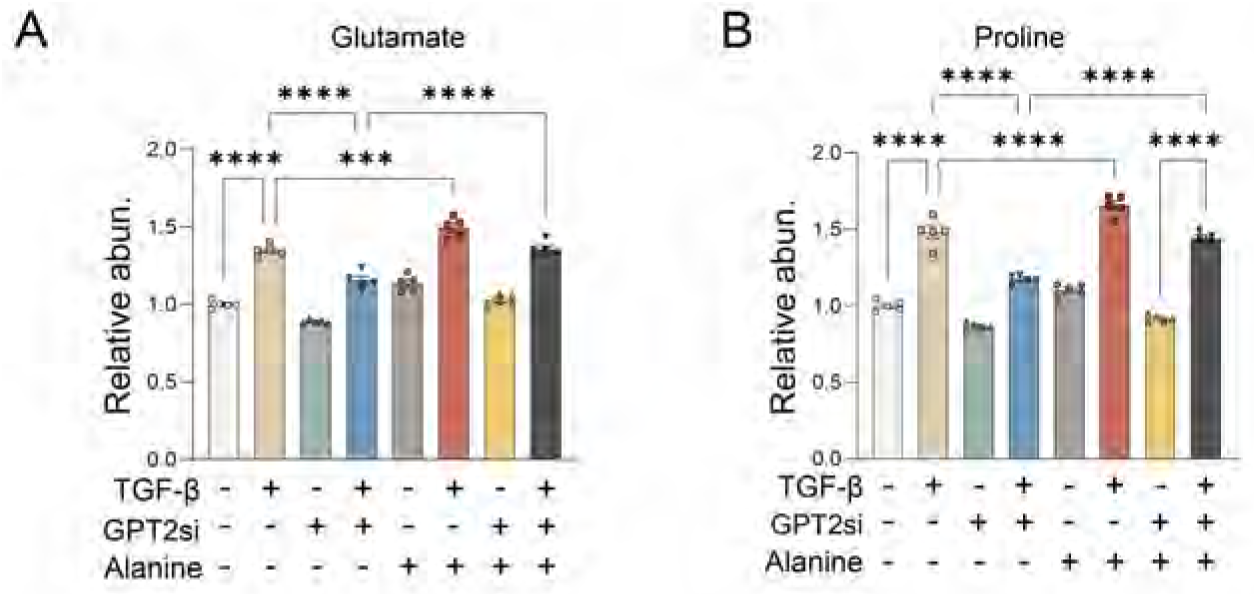
Alanine deprivation disrupts glutamate and proline metabolic pathways. (A, B) Relative quantification of intracellular glutamate (A), and proline (B) levels in GPT2 knockdown NHLFs cultured in DMEM with TGF-β (48 h) and supplemented with alanine (2 mM) (n = 5 per group). A and B: One-way ANOVA. Data are presented as mean ± SEM. ***p < 0.001; ****p < 0.0001.

**Figure EV7.**
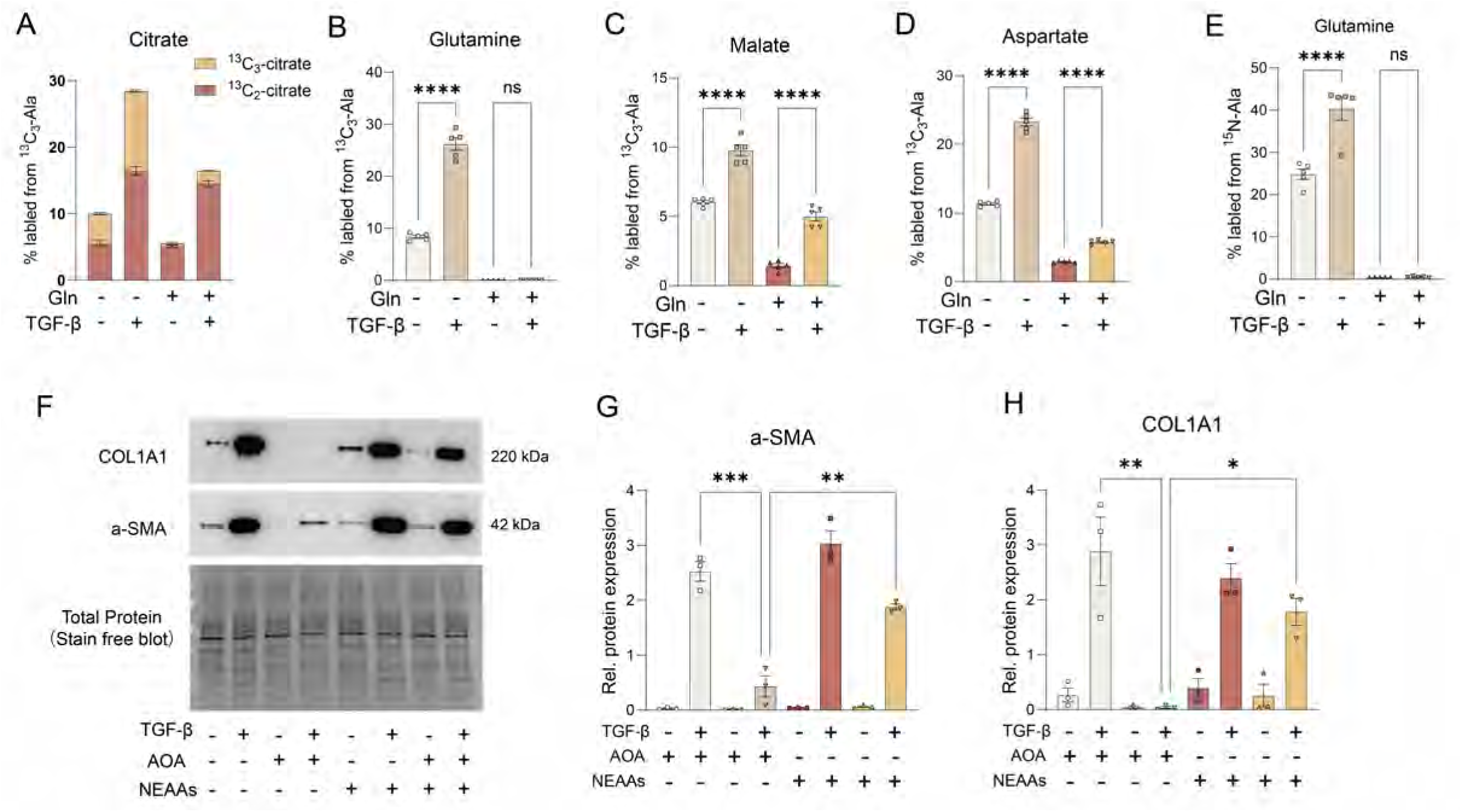
Alanine is a carbon and nitrogen source for anabolic metabolism during myofibroblast differentiation. (A-D) Metabolite labeling from [U-^13^C_3_]-alanine showing fractional label enrichment in intracellular ^13^C_2_- or ^13^C_3_-citrate (A), glutamine (B), malate (C), and aspartate(D), with or without TGF-β treatment in the presence or absence of exogenous glutamine. (E) Fractional label enrichment of intracellular glutamine from ^15^N-alanine. (F) Representative Western blot showing α-SMA and COL1A1 expression in NHLFs cultured in DMEM with TGF-β (2 ng/mL) for 48 h, in the presence or absence of the GPT inhibitor AOA (1 mM) and/or NEAA cocktail supplementation. (G, H) Quantification of α-SMA (F) and COL1A1 (G) protein levels relative to total protein. A-E, G, and H: One-way ANOVA. Data are presented as mean ± SEM. ns, p > 0.05; *p < 0.05; **p < 0.01; ***p < 0.001; ****p < 0.0001.

**Figure EV8.**
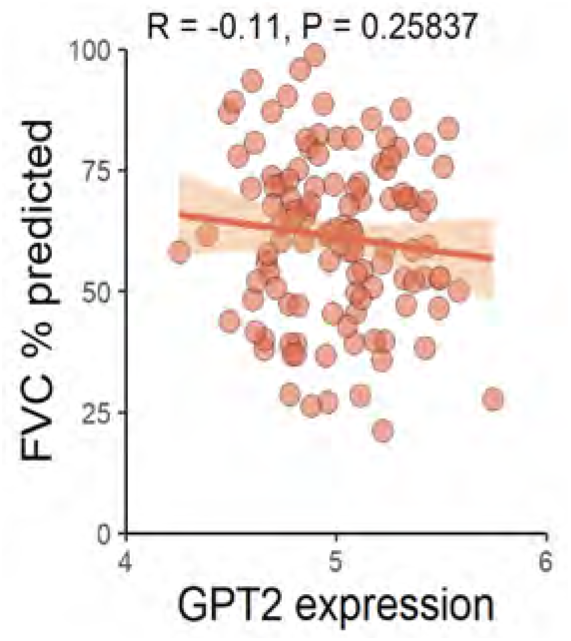
Correlation between GPT2 expression and pulmonary function. Pearson’s correlation between GPT2 mRNA expression and predicted forced vital capacity (FVC) in patients, based on clinical and transcriptomic data from GSE32537. No significant correlation was observed (R = –0.11, P = 0.258).

